# An active inference approach to modeling structure learning: concept learning as an example case

**DOI:** 10.1101/633677

**Authors:** Ryan Smith, Philipp Schwartenbeck, Thomas Parr, Karl J. Friston

## Abstract

Within computational neuroscience, the algorithmic and neural basis of structure learning remains poorly understood. Concept learning is one primary example, which requires both a type of internal model expansion process (adding novel hidden states that explain new observations), and a model reduction process (merging different states into one underlying cause and thus reducing model complexity via meta-learning). Although various algorithmic models of concept learning have been proposed within machine learning and cognitive science, many are limited to various degrees by an inability to generalize, the need for very large amounts of training data, and/or insufficiently established biological plausibility. Using concept learning as an example case, we introduce a novel approach for modeling structure learning – and specifically state-space expansion and reduction – within the active inference framework and its accompanying neural process theory. Our aim is to demonstrate its potential to facilitate a novel line of active inference research in this area. The approach we lay out is based on the idea that a generative model can be equipped with extra (hidden state or cause) ‘slots’ that can be engaged when an agent learns about novel concepts. This can be combined with a Bayesian model reduction process, in which any concept learning – associated with these slots – can be reset in favor of a simpler model with higher model evidence. We use simulations to illustrate this model’s ability to add new concepts to its state space (with relatively few observations) and increase the granularity of the concepts it currently possesses. We also simulate the predicted neural basis of these processes. We further show that it can accomplish a simple form of ‘one-shot’ generalization to new stimuli. Although deliberately simple, these simulation results highlight ways in which active inference could offer useful resources in developing neurocomputational models of structure learning. They provide a template for how future active inference research could apply this approach to real-world structure learning problems and assess the added utility it may offer.

## Introduction

The ability to learn the latent structure of one’s environment – such as inferring the existence of hidden causes of regularly observed patterns in co-occurring feature observations – is central to human cognition. For example, we do not simply observe particular sets of colors, textures, shapes, and sizes – we also observe *identifiable object*s such as, say, a ‘screwdriver’. If we were tool experts, we might also recognize particular types of screwdrivers (e.g., flat vs. Phillip’s head), designed for a particular use. This ability to learn latent structure, such as learning to recognize co-occurring features under conceptual categories (as opposed to just perceiving sensory qualities; e.g., red, round, etc.), is also highly adaptive. Only if we knew an object was a screwdriver could we efficiently infer that it affords putting certain structures together and taking them apart; and only if we knew the specific type of screwdriver could we efficiently infer, say, the artefacts to use it on. Many concepts of this sort require experience-dependent acquisition (i.e., learning).

From a computational perspective, the ability to acquire a new concept can be seen as a type of structure learning involving Bayesian model comparison (MacKay and Peto, 1995; Botvinick et al., 2009; Gershman and Niv, 2010; Salakhutdinov et al., 2013a; Tervo et al., 2016). Specifically, concept acquisition can be cast as an agent learning (or inferring) that a new hypothesis (referred to here as a hidden cause or state) should be added to the internal or generative model with which it explains its environment, because existing causes cannot account for new observations (e.g., an agent might start out believing that the only tools are hammers and screwdrivers, but later learn that there are also wrenches). In other words, the structure of the space of hidden causes itself needs to expand to accommodate new patterns of observations. This model expansion process is complementary to a process called Bayesian model reduction (Friston and Penny, 2011); in which the agent can infer that there is redundancy in its model, and a model with fewer states or parameters provides a more parsimonious (i.e. simpler) explanation of observations (Schmidhuber, 2006; Friston et al., 2017b). For example, in some instances it may be more appropriate to differentiate between fish and birds as opposed to salmon, peacocks and pigeons. This reflects a reduction in model complexity based on a particular feature space underlying observations and thus resonates with other accounts of concept learning as dimensionality reduction (Stachenfeld et al., 2016; Behrens et al., 2018) – a topic we discuss further below.

A growing body of work in a number of domains has approached this problem from different angles. In developmental psychology and cognitive science, for example, probability theoretic (Bayesian) models have been proposed to account for word learning in children and the remarkable human ability to generalize from very few (or even one) examples in which both broader and narrower categorical referents could be inferred (Kemp et al., 2007; Xu and Tenenbaum, 2007a, 2007b; Perfors et al., 2011; Lake et al., 2015). In statistics, a number of nonparametric Bayesian models, such as the “Chinese Room” process and the “Indian Buffet” process, have been used to infer the need for model expansion (Gershman and Blei, 2012). There are also related approaches in machine learning, as applied to things like Gaussian mixture models (McNicholas, 2016), as well as models based on Bayesian program learning (Lake et al., 2015) and combinations of deep learning and hierarchical Bayesian methods (Salakhutdinov et al., 2013b).

Such approaches often employ clustering algorithms, which take sets of data points in a multidimensional space and divide them into separable clusters (e.g., see (Anderson, 1991; Love et al., 2004; Sanborn et al., 2010)). While many of these approaches assume the number of clusters is known in advance, various goodness-of-fit criteria may be used to determine the optimal number. However, a number of approaches require much larger amounts of data than humans do to learn new concepts (Geman et al., 1992; Lecun et al., 1998; Hinton et al., 2012; LeCun et al., 2015; Mnih et al., 2015). Many also require corrective feedback to learn and yet fail to acquire sufficiently rich conceptual structure to allow for generalization (Osherson and Smith, 1981; Barsalou, 1983; Biederman, 1987; Ward, 1994; Feldman, 1997; Markman and Makin, 1998; Williams and Lombrozo, 2010; Jern and Kemp, 2013).

Approaches to formally modeling structure learning, including concept learning, have not yet been examined within the emerging field of research on Active Inference models within computational neuroscience (Friston, 2010; Friston et al., 2016a, 2017b, 2017c). This represents a novel and potentially fruitful research avenue that, as discussed below, may offer unique advantages in research focused on understanding the neural basis of learning latent structure. In this paper, we explore the potential of this approach. We aim to provide a type of modeling template that could be used in future active inference research on real-world structure learning problems – and assess the additional utility it might offer. We provide a number of example simulations demonstrating how structure learning can be seen as an emergent property of active inference (and learning) under generative models with ‘spare capacity’; where spare or uncommitted capacity is used to expand the repertoire of representations (Baker and Tenenbaum, 2014), while Bayesian model reduction (Hobson and Friston, 2012; Friston et al., 2017b) promotes generalization by minimizing model complexity – and releasing representations to replenish ‘spare capacity’.

From a machine learning perspective, Bayesian model reduction affords the opportunity to consider generative models with a large number of hidden states or latent factors and then optimize the number (or indeed partitions) of latent factors with respect to a variational bound on model evidence. This could be regarded as a bounded form of nonparametric Bayes, in which a potentially infinite number of latent factors are considered; with appropriate (e.g., Indian buffet process) priors over the number of hidden states generating data features^1^. The approach we articulate here is bounded in the sense that an upper bound on the number of hidden states is specified prior to structure learning. Furthermore, in virtue of the (biologically plausible) variational schemes used for model reduction, there is no need to explicitly compute model evidence; thereby emulating the computational efficiency of nonparametric Bayes (Gershman and Blei, 2012), while accommodating any prior over models.

In what follows, we first provide a brief overview of active inference. We then introduce a model of concept learning (using basic and subordinate level animal categories), as a representative example of structure learning. We specifically model cognitive (semantic) processes that add new concepts to a state space and that optimize the granularity of an existing state space. We then establish the validity of this model using numerical analyses of concept learning, when repeatedly presenting a synthetic agent with different animals characterized by different combinations of observable features. We demonstrate how particular approaches combining Bayesian model reduction and expansion can reproduce successful concept learning without the need for corrective feedback – and allow for generalization. We further demonstrate the ability of this model to generate predictions about measurable neural responses based on the neural process theory that accompanies active inference. We conclude with a brief discussion of the implications of this work. Our goal is to present an introductory proof of concept – that could be used as the foundation of future active inference research on the neurocomputational basis of structure learning.

### An Active Inference approach for modeling concept learning through state-space expansion

#### A primer on Active Inference

Active Inference suggests that the brain is an inference machine that approximates optimal probabilistic (Bayesian) belief updating across perceptual, cognitive, and motor domains. Active Inference more specifically postulates that the brain embodies an internal model of the world that is “generative” in the sense that it can simulate the sensory data that it should receive if its model of the world is correct. These simulated (predicted) sensory data can be compared to actual observations, and differences between predicted and observed sensations can be used to update the model. Over short timescales (e.g., a single observation) this updating corresponds to inference (perception), whereas on longer timescales it corresponds to learning (i.e., updating expectations about what will be observed later). Another way of putting this is that perception optimizes beliefs about the current state of the world, while learning optimizes beliefs about the relationships between the variables that constitute the world. These processes can be seen as ensuring the generative model (entailed by recognition processes in the brain) remains an accurate model of the world that it seeks to regulate (Conant and Ashbey, 1970).

Active Inference casts decision-making in similar terms. Actions can be chosen to resolve uncertainty about variables within a generative model (i.e., sampling from domains in which the model does not make precise predictions), which can prevent anticipated deviations from predicted outcomes. In addition, some expectations are treated as a fixed phenotype of an organism. For example, if an organism did not continue to “expect” to observe certain amounts of food, water, and shelter, then it would quickly cease to exist (McKay and Dennett, 2009) – as it would not pursue those behaviors that fulfill these expectations (c.f. the ‘optimism bias’ (Sharot, 2011)). Thus, a creature should continually seek out observations that support – or are internally consistent with – its own continued existence. Decision-making can therefore be cast as a process in which the brain infers the sets of actions (policies) that would lead to observations consistent with its own survival-related expectations (i.e., its “prior preferences”). Mathematically, this can be described as selecting sequences of actions (policies) that maximize “Bayesian model evidence” expected under a policy, where model evidence is the (marginal) likelihood that particular sensory inputs would be observed under a given model.

In real-world settings, directly computing Bayesian model evidence is generally intractable. Thus, some approximation is necessary. Active Inference proposes that the brain computes a quantity called “variational free energy” that provides a bound on model evidence, such that minimization of free energy indirectly maximizes model evidence (this is exactly the same functional used in machine learning where it is known as an evidence lower bound or ELBO). In this case, decision-making will be approximately (Bayes) optimal if it infers (and enacts) the policy that will minimize expected free energy (i.e., free energy with respect to a policy, where one takes expected future observations into account). Technically, expected free energy is the average free energy under the posterior predictive density over policy-specific outcomes.

Expected free energy can be decomposed in different ways that reflect uncertainty and prior preferences, respectively (e.g., epistemic and instrumental affordance or ambiguity and risk). This formulation means that any agent that minimizes expected free energy engages initially in exploratory behavior to minimize uncertainty in a new environment. Once uncertainty is resolved, the agent then exploits that environment to fulfill its prior preferences. The formal basis for Active Inference has been thoroughly detailed elsewhere (Friston et al., 2017a), and the reader is referred there for a full mathematical treatment.

When the generative model is formulated as a partially observable Markov decision process (a mathematical framework for modeling decision-making in cases where some outcomes are under the control of the agent and others are not, and where states of the world are not directly known but must be inferred from observations), active inference takes a particular form. Here, the generative model is specified by writing down plausible or allowable policies, hidden states of the world (that must be inferred from observations), and observable outcomes, as well as a number of matrices that define the probabilistic relationships between these quantities (see right panel of figure 1).

**Figure 1.**
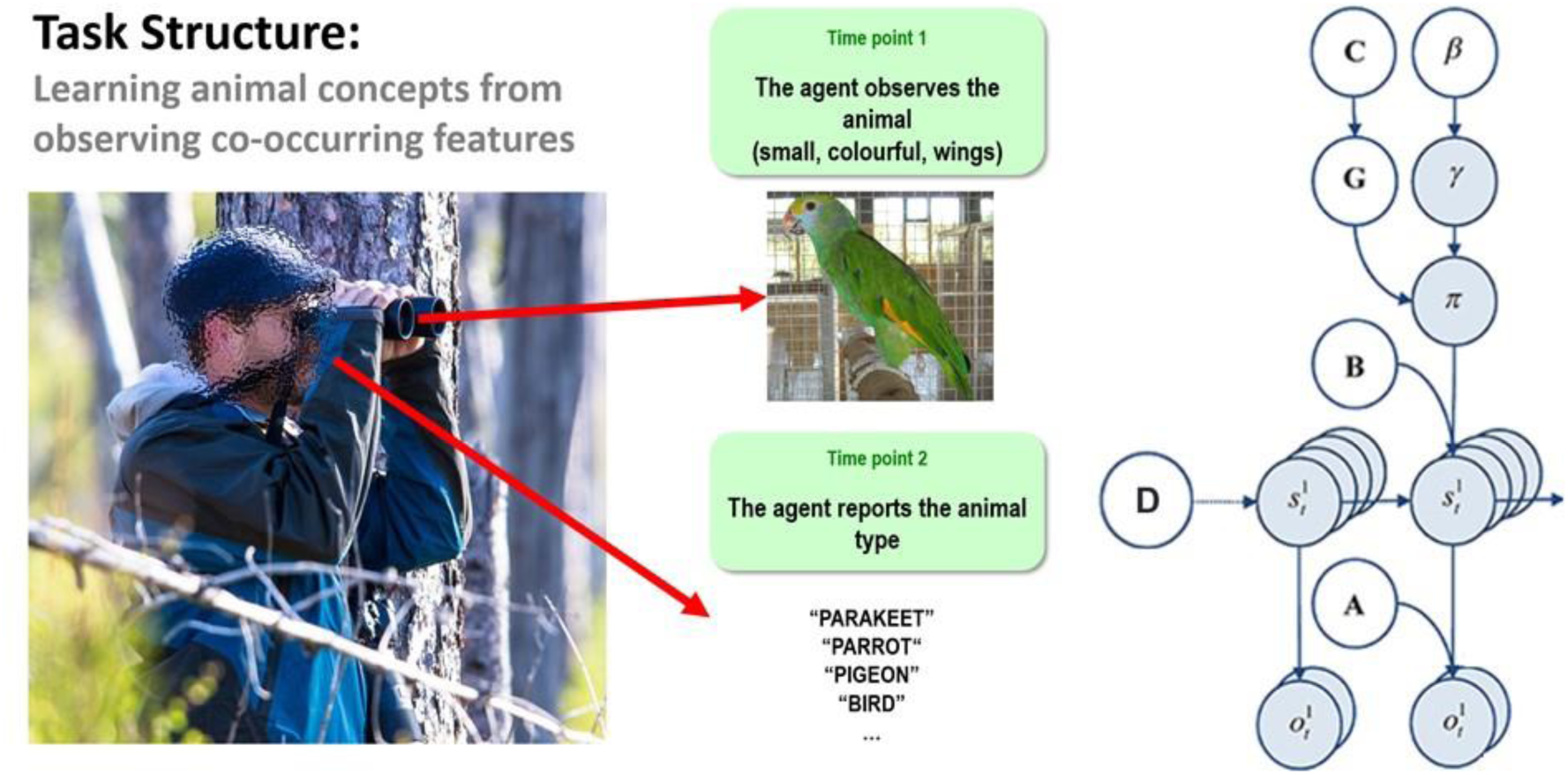
Left: Illustration of the trial structure performed by the agent. At the first time point, the agent is exposed to one of 8 possible animals that are each characterized by a unique combination of visual features. At the 2^nd^ time point, the agent would then report which animal concept matched that feature combination. The agent could report a specific category (e.g., pigeon, hawk, minnow, etc.) or a general category (i.e., bird or fish) if insufficiently certain about the specific category. See the main text for more details. Right: Illustration of the Markov decision process formulation of active inference used in the simulations described in this paper. The generative model is here depicted graphically, such that arrows indicate dependencies between variables. Here observations (**o**) depend on hidden states (**s**), as specified by the **A** matrix, and those states depend on both previous states (as specified by the **B** matrix, or the initial states specified by the **D** matrix) and the policies (**π**) selected by the agent. The probability of selecting a particular policy in turn depends on the expected free energy (**G**) of each policy with respect to the prior preferences (**C**) of the agent. The degree to which expected free energy influences policy selection is also modulated by a prior policy precision parameter (**γ**), which is in turn dependent on beta (***β***) –where higher values of beta promote more randomness in policy selection (i.e., less influence of the differences in expected free energy across policies). For more details regarding the associated mathematics, see (KJ Friston, Lin, et al., 2017; KJ Friston, Parr, et al., 2017).

The ‘**A**’ matrix indicates which observations are generated by each combination of hidden states (i.e., the likelihood mapping specifying the probability that a particular set of observations would be observed given a particular set of hidden states). The ‘**B**’ matrix is a transition matrix, indicating the probability that one hidden state will evolve into another over time. The agent, based on the selected policy, controls some of these transitions (e.g., those that pertain to the positions of its limbs). The ‘**D**’ matrix encodes prior expectations about the initial hidden state the agent will occupy. Finally, the ‘**C**’ matrix specifies prior preferences over observations; it quantifies the degree to which different observed outcomes are rewarding or punishing to the agent. In these models, observations and hidden states can be factorized into multiple outcome *modalities* and hidden state *factors*.

This means that the likelihood mapping (the **A** matrix) can also model the interactions among different hidden states when generating outcomes (observations).

### From principles to process theories

One potential empirical advantage of the present approach stems from the fact that active inference models have a plausible biological basis that affords testable neurobiological predictions. Specifically, these models have well-articulated companion micro-anatomical neural process theories, based on commonly used message-passing algorithms (Friston et al., 2017a; Parr and Friston, 2018; Parr et al., 2019). In these process theories, for example, the activation level of different neural populations (typically portrayed as consisting of different cortical columns) can encode posterior probability estimates over different hidden states. These activation levels can then be updated by synaptic inputs with particular weights that convey the conditional probabilities encoded in the **A** and **B** (among other) matrices described above, where active learning then corresponds to associative synaptic plasticity. Phasic dopamine responses also play a particular role in these models, by reporting changes in policy precision (i.e., the degree of confidence in one policy over others) upon new observations (see Figure 2 and the associated legend for more details). In what follows, we describe how the type of generative model – that underwrites these processes – was specified to perform concept inference/learning.

**Figure 2.**
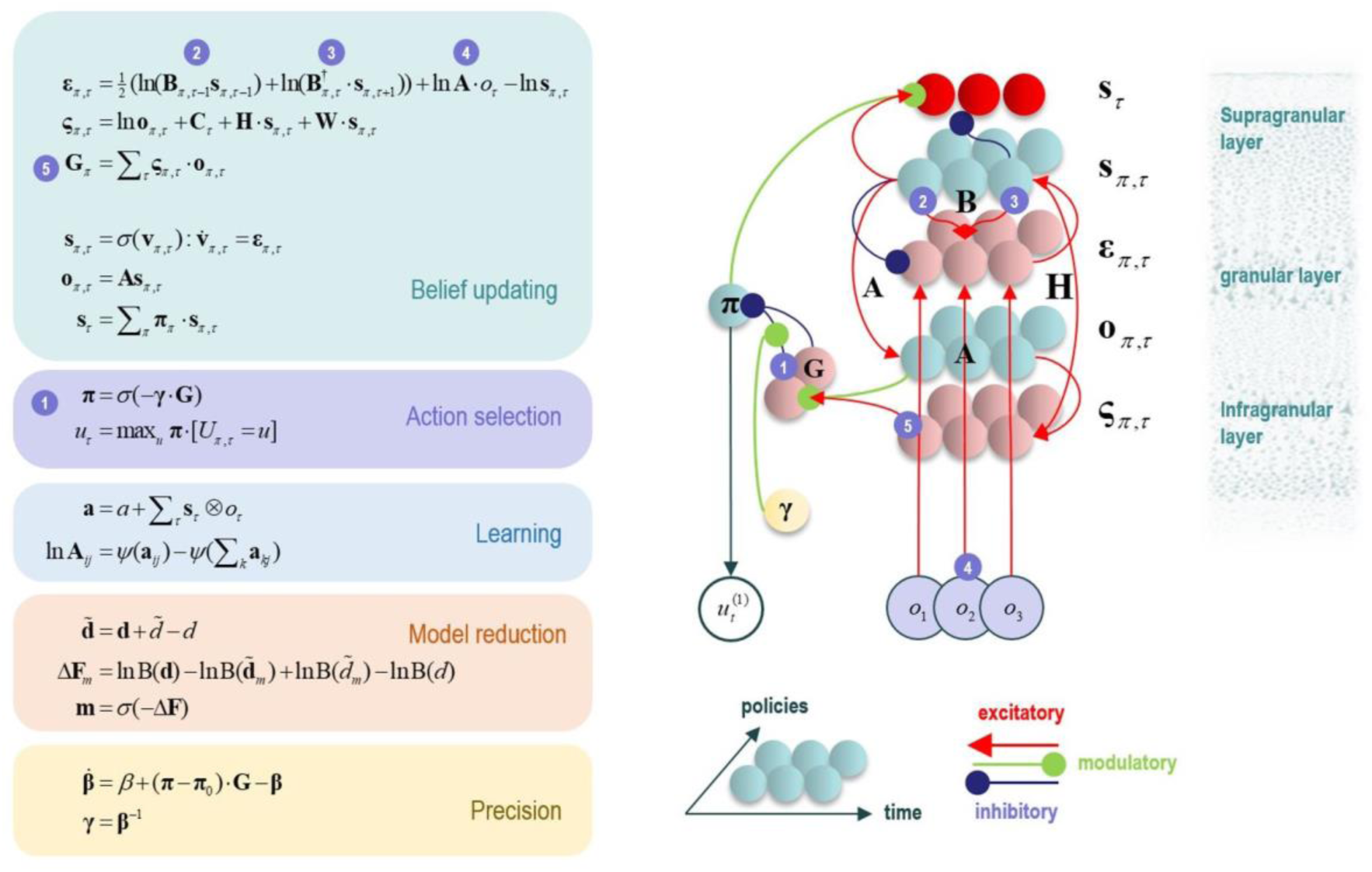
This figure illustrates the mathematical framework of active inference and associated neural process theory used in the simulations described in this paper. The differential equations in the left panel approximate Bayesian belief updating within the graphical model depicted in the right panel of Figure 1, via a gradient descent on free energy (**F**). The right panel illustrates a possible neural basis by which neurons making up cortical columns could implement these equations. The equations have been expressed in terms of two types of prediction errors. State prediction errors (**ε**) are used to update expectations over states (**s**) by driving depolarization (**v**) in neurons encoding those hidden states. The probability distribution over hidden states is then obtained via a softmax (normalized exponential) function (**σ**). Outcome prediction errors (**ς**) convey the difference between preferred observations and those predicted under each policy (**π**), and are used to evaluate the expected free energy (**G**) of each policy. This term additionally considers the expected ambiguity or conditional entropy (**H**) between states and outcomes, as well as a novelty term (**W**) reflecting the degree to which beliefs about how states generate outcomes would change upon observing different possible state-outcome mappings. Outcome prediction errors thus allow policies to be evaluated based on both expected information gain and expected reward. Policy-specific free energies are then integrated to give the policy probabilities via a softmax function. Actions at each time point (**u**) are then chosen out of the possible actions under each policy (**U**), weighted by the probability of each policy. In our simulations, the model learned associations between hidden states and observations (**A**) via a process in which counts were accumulated (**a**) reflecting the number of times the agent observed a particular outcome when it believed that it occupied each possible hidden state. Although not displayed explicitly, learning prior expectations over initial hidden states (**D**) is similarly accomplished via accumulation of concentration parameters (**d**). These prior expectations reflect counts of how many times the agent believes it previously occupied each possible initial state. Concentration parameters are converted into expected log probabilities using digamma functions (***ψ***). The way in which Bayesian model reduction was performed in this paper is also written in the lower left (where B indicates a beta function, and **m** is the posterior probability of each model). Here, the posterior distribution over initial states (**d**) is used to assess the difference in the evidence (**ΔF**) it provides for the number of hidden states in the current model and other possible models characterized by fewer hidden states. Prior concentration parameters are shown in italics, posteriors in bold, and those priors and posteriors associated with the reduced model are equipped with a tilde (**∼**). As already stated, the right panel illustrates a possible neural implementation of the update equations in the left panel. In this implementation, probability estimates have been associated with neuronal populations that are arranged to reproduce known intrinsic (within cortical area) connections. Red connections are excitatory, blue connections are inhibitory, and green connections are modulatory (i.e., involve a multiplication or weighting). These connections mediate the message passing associated with the equations in the left panel. Note that the various messages passed between neurons to implement the equations on the left are denoted with corresponding numbers in the left and right panels – conveying policy values (message 1), prior expectations (messages 2,3), sensory signals (message 4), and expected free energies (message 5). Cyan units correspond to expectations about hidden states and (future) outcomes under each policy, while red states indicate their Bayesian model averages (i.e., a “best guess” based on the average of the probability estimates for the states and outcomes across policies, weighted by the probability estimates for their associated policies). Pink units correspond to (state and outcome) prediction errors that are averaged to evaluate expected free energy and subsequent policy expectations (in the lower part of the network). This (neural) network formulation of belief updating means that connection strengths correspond to the parameters of the generative model described in the text. Learning then corresponds to changes in the synaptic connection strengths. Only exemplar connections are shown to avoid visual clutter. Furthermore, we have just shown neuronal populations encoding hidden states under two policies over three time points (i.e., two transitions), whereas in the task described in this paper there are greater number of allowable policies. For more information regarding the mathematics and processes illustrated in this figure, see (KJ Friston, Lin, et al., 2017; KJ Friston, Parr, et al., 2017).

### A model of concept inference and learning through state-space expansion

To model concept inference, we constructed a simple task for an agent to perform (see figure 1, left panel). In this task, the agent was presented with different animals on different trials and asked to answer a question about the type of animal that was seen. As described below, in some simulations the agent was asked to report the type of animal that was learned previously; in other simulations, the agent was instead asked a question that required conceptual generalization.

Crucially, to answer these questions the agent was required to observe different animal features, where the identity of the animal depended on the combination of features. There were three feature categories (size, color, and species-specific; described further below) and two discrete time points in a trial (observe and report).

To simulate concept learning and state-space expansion (based on the task described above) we needed to specify an appropriate generative model. Once this model has been specified, one can use standard (variational) message passing to simulate belief updating and behavior in a biologically plausible way: for details, please see (Friston et al., 2017a, 2017c). In our (minimal) model, the first hidden state factor included (up to) eight levels, specifying four possible types of birds and four possible types of fish (Figure 3A). The outcome modalities included: a feature space including two size features (big, small), two color features (gray, colorful), and two species-differentiating features (wings, gills). The **A** matrix specified a likelihood mapping between features and animal concepts, such that each feature combination was predicted by an animal concept (Hawk, Pigeon, Parrot, Parakeet, Sturgeon, Minnow, Whale shark, Clownfish). This model was deliberately simple to allow for a clear illustration, but it is plausibly scalable to include more concepts and a much larger feature space. The **B** matrix for the first hidden state factor was an identity matrix, reflecting the belief that the animal identity was conserved during each trial (i.e., the animals were not switched out mid-trial).

**Figure 3.**
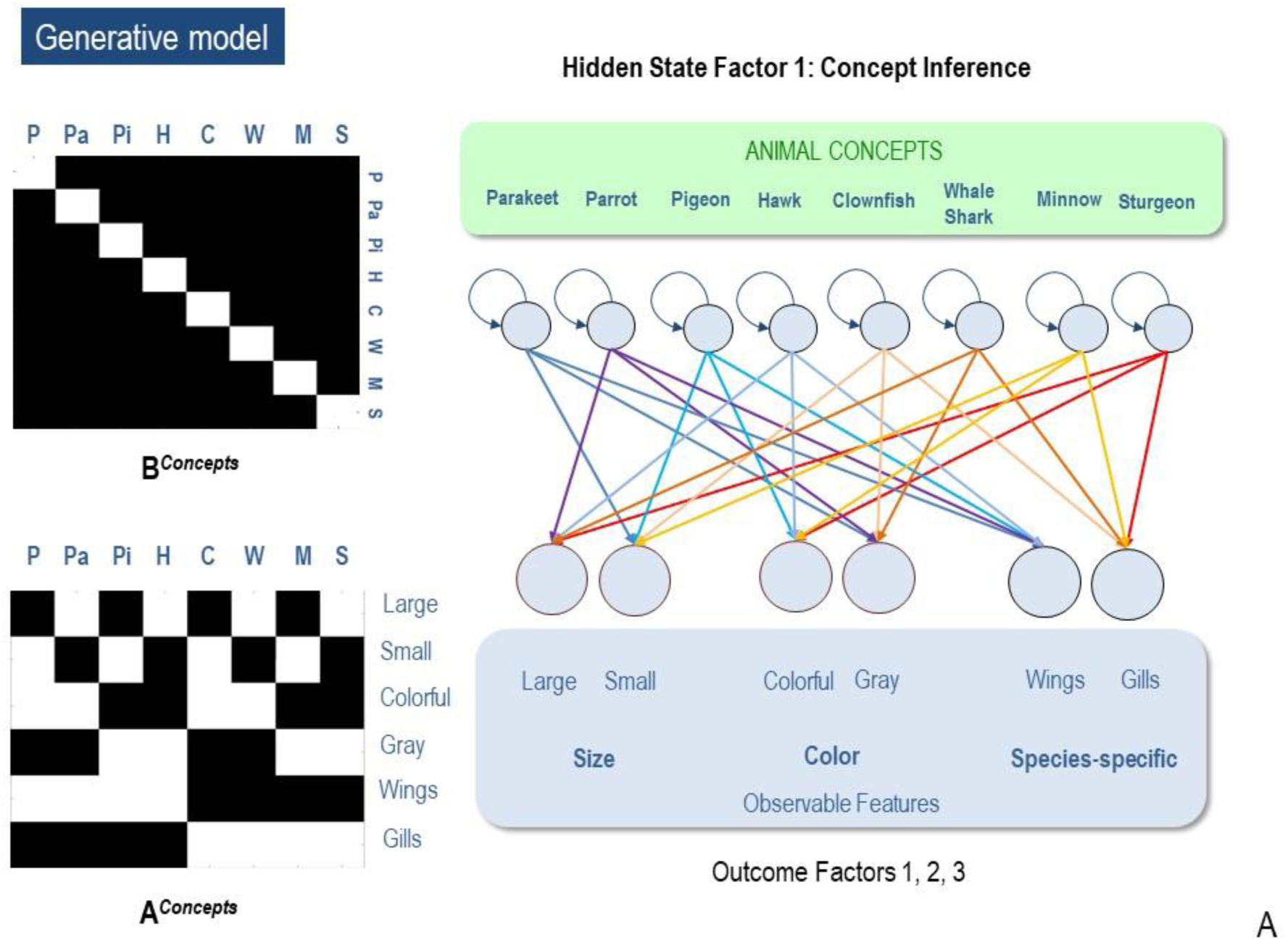
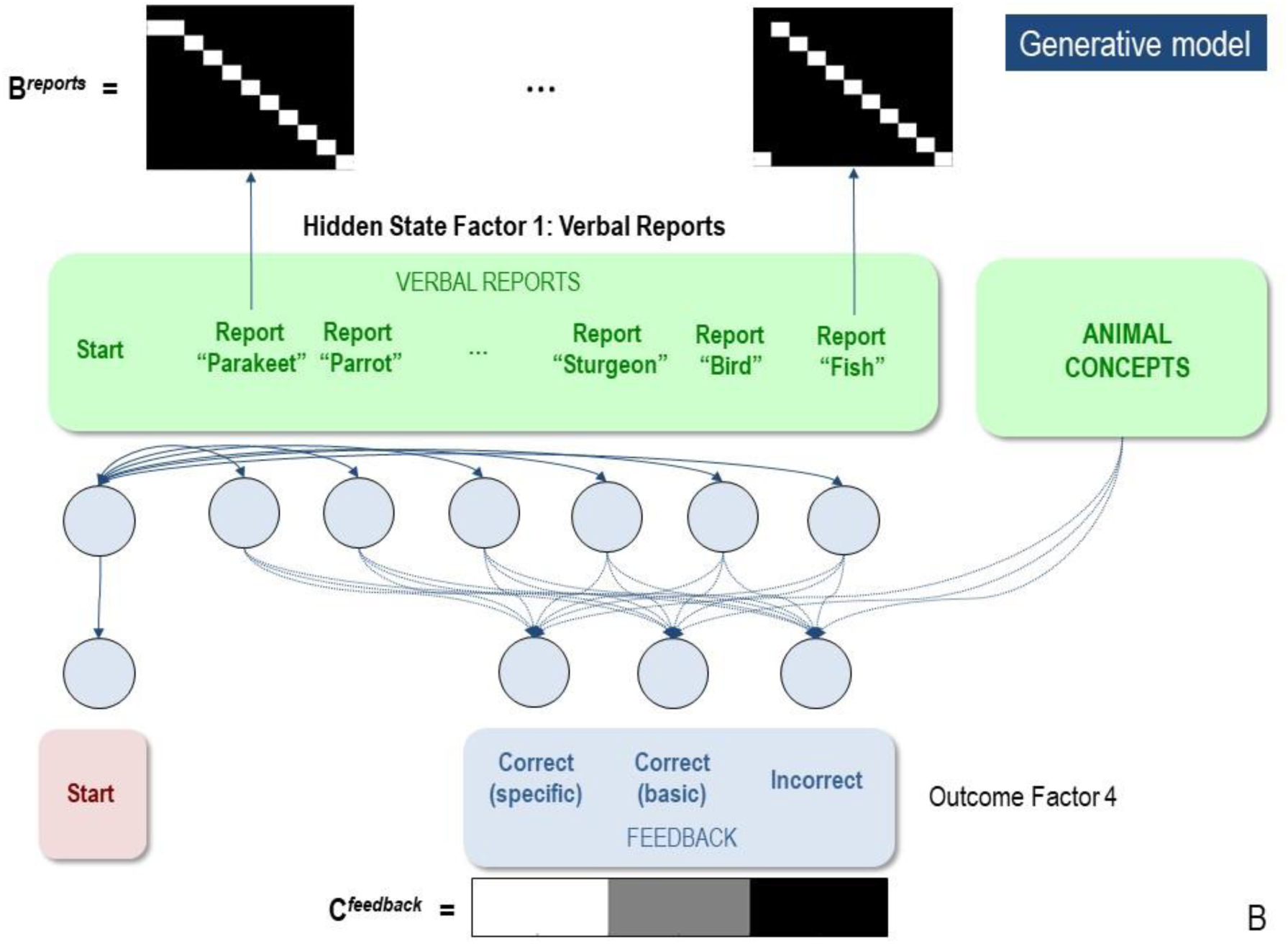
(A) Illustration of the first hidden state factor containing columns (levels) for 8 different animal concepts. Each of these 8 concepts generated a different pattern of visual feature observations associated with the outcome modalities of size, color, and species-specific features. The **B** matrix was an identity matrix, indicating that the animal being observed did not change within a trial (white = 1, black = 0). The **A** matrix illustrates the specific mapping from animal concepts to feature combinations. As depicted, each concept corresponded to a unique point in a 3-dimensional feature space. (B) illustration of the 2^nd^ hidden state factor corresponding to the verbal reports the agent could choose in response to its observations. These generated feedback as to whether its verbal report was accurate with respect to a basic category report or a specific category report. As illustrated in the **C** matrix, the agent most preferred to be correct about specific categories, but least preferred being incorrect. Thus, reporting the basic categories was a safer choice if the agent was too uncertain about the specific identity of the animal.

The second hidden state factor was the agent’s report. That this is assumed to factorize from the first hidden state factor means that there is no prior constraint that links the chosen report to the animal generating observations. The agent could report each of the eight possible specific animal categories, or opt for a less specific report of a bird or a fish. Only one category could be reported at any time. Thus, the agent had to choose to report only bird vs. fish or to report a more specific category. In other words, the agent could decide upon the appropriate level of coarse-graining of its responses (figure 3B).

During learning trials, the policy space was restricted such that the agent could not provide verbal reports or observe corrective feedback (i.e., all it could do is “stay still” in its initial state and observe the feature patterns presented). This allowed the agent to learn concepts in an unsupervised manner (i.e. without being told what the true state was or whether it was correct or incorrect). After learning, active reporting was enabled, and the **C** matrix was set so that the agent preferred to report correct beliefs. We defined the preferences of the agent such that it preferred correctly reporting specific category knowledge and was averse to incorrect reports. This ensured that it only reported the general category of bird vs. fish, unless sufficiently certain about the more specific category.

In the simulations reported below, there were two time points in each trial of categorisation or conceptual inference. At the first time point, the agent was presented with the animals features, and always began in a state of having made no report (the “start” state). The agent’s task was simply to observe the features, infer the animal identity, and then report it (i.e., in reporting trials). Over 32 simulations (i.e., 4 trials per animal), we confirmed that, if the agent already started out with full knowledge of the animal concepts (i.e., a fully precise **A** matrix), it would report the specific category correctly 100% of the time. Over an additional 32 simulations, we also confirmed that, if the agent was only equipped with knowledge of the distinction between wings and gills (i.e., by replacing the rows in the **A** matrix corresponding to the mappings from animals to size and color with flat distributions), it would report the generic category correctly 100% of the time but would not report the specific categories.^2^ This was an expected and straightforward consequence of the generative model – but provides a useful example of how agents trade off preferences and different types of uncertainty.

### Simulating concept learning and the acquisition of expertise

Having confirmed that our model could successfully recognize animals if equipped with the relevant concepts (i.e., likelihood mappings) – we turn now to concept learning.

### Concept acquisition through state-space expansion

We first examined our model’s ability to acquire concept knowledge in two distinct ways. This included 1) the agent’s ability to effectively “expand” (i.e., fill in an unused column within) its state space and add new concepts and 2) the agent’s ability to increase the granularity of its conceptual state space and learn more specific concepts, when it already possessed broader concepts.

#### Adding Concepts

To assess whether our agent could effectively expand its state space and acquire a new concept, we first set one column of the previously described model’s **A** matrix (mapping an animal concept to its associated features) to be a uniform distribution^3^; creating an imprecise likelihood mapping for one concept – essentially, that concept predicted all features with nearly equal probability. Here, we chose sturgeon (large, gray, gills) as the concept for which the agent had no initial knowledge (see Figure 4A, right-most column of left-most ‘pre-learning’ matrix). We then generated 2000 observations based on the outcome statistics of a model with full knowledge of all eight animals (the “generative process”), to test whether the model could infer that a novel animal was present and then learn the correct likelihood mapping for sturgeon (note: this excessive number of observations was used for consistency with later simulations, in which more concepts had to be learned, and also to evaluate how performance improved as a function of the number of observations the agent was exposed to; see figure 4B).

**Figure 4.**
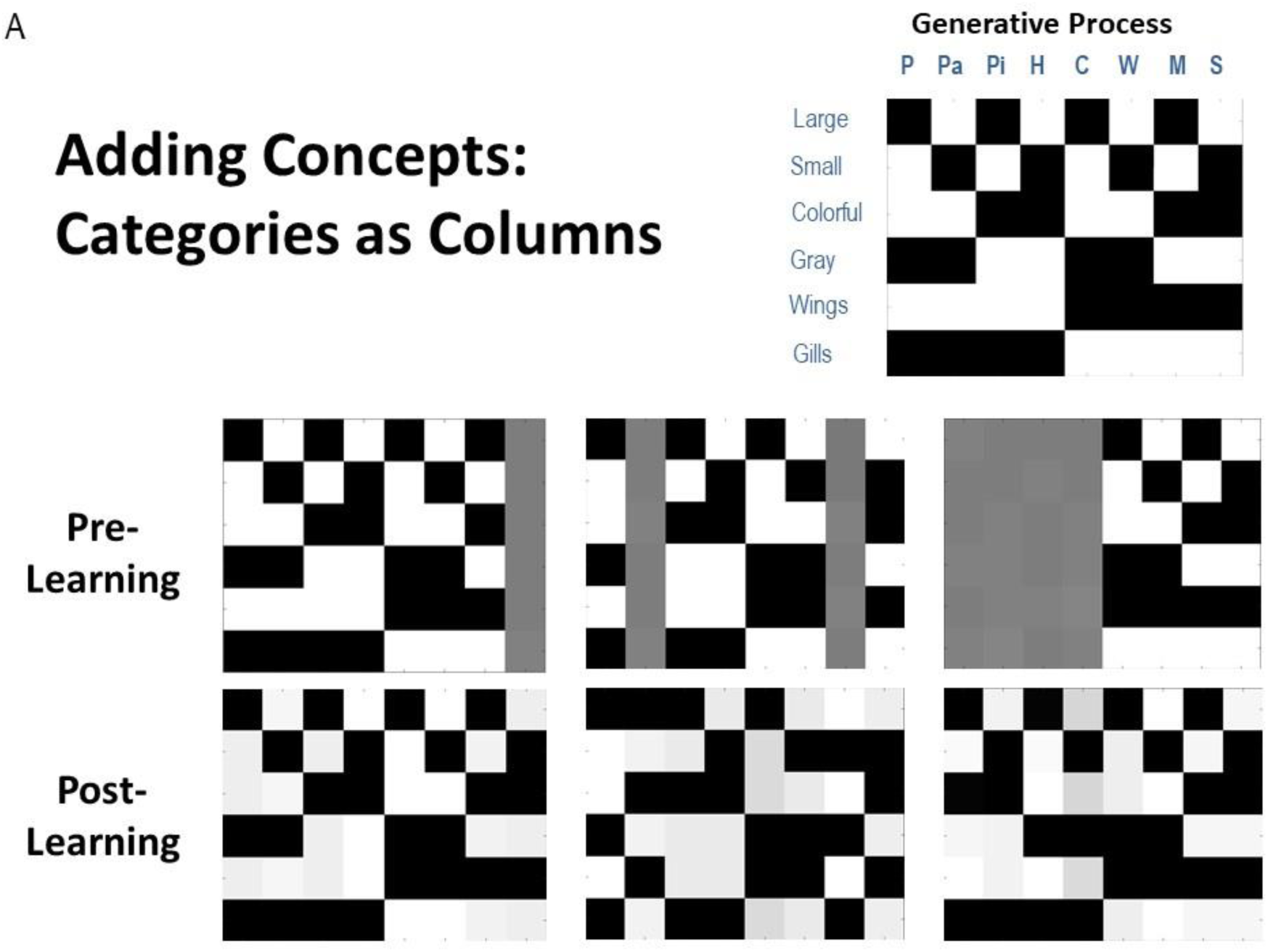
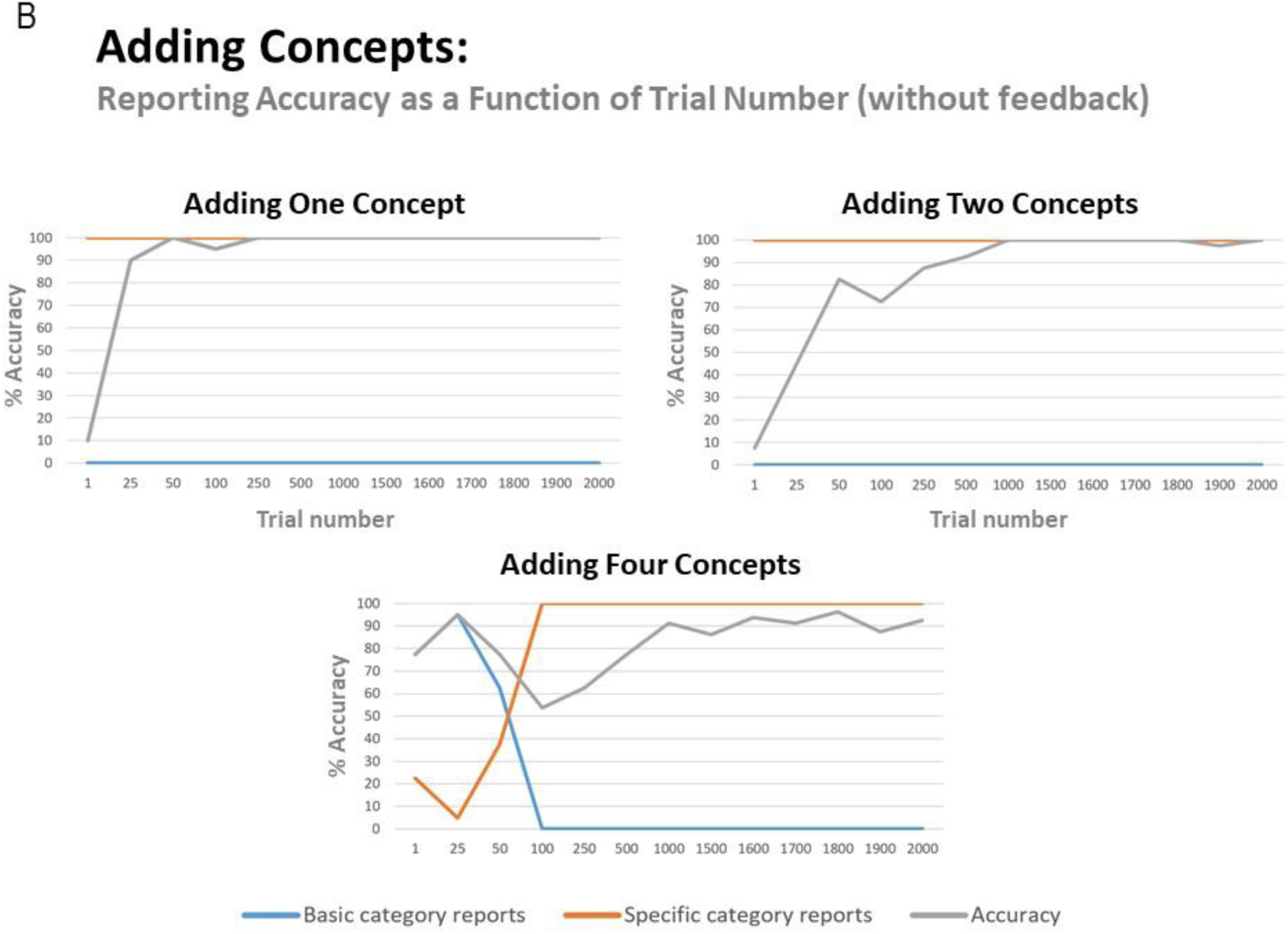
(A) illustration of representative simulation results in which the agent successfully learned 1, 2, or 4 new animal concept categories with no prior knowledge beforehand. The generative process is shown in the upper right, illustrating the feature combinations to be learned. Pre-learning, either 1, 2 or 4 columns in the likelihood mapping began as a flat distribution with a slight amount of Gaussian noise. The agent was then provided with 2000 observations of the 8 animals with equal probability. Crucially, the agent was prevented from providing verbal reports during these 2000 trials and thus did not receive feedback about the true identity of the animal. Thus, learning was driven completely by repeated exposure in an unsupervised manner. Also note that, while the agent was successful at learning the new concepts, it did not always assign the new feature patterns to the same columns as illustrated in the generative process. This is to be expected given that the agent received no feedback about the true hidden state that generated its observations. (B) illustration of how reporting accuracy, and the proportion of basic category and specific category responses, changed as a function of repeated exposures. This was accomplished by taking the generative model at a number of representative trials and then testing it with 20 observations of each animal in which reporting was enabled. As can be seen, maximal accuracy was achieved much more quickly when the agent had to learn fewer concepts. When it had learned 4 concepts, it also began by reporting the general categories and then subsequently became sufficiently confident to report the more specific categories.

We refer to this as “effective” state space expansion because the dimensions of the hidden state space do not change. Instead, based on the model structure described above, the agent begins with implicit prior expectations about the structure of its environment (‘structural priors’) – namely, that there could be up to 8 different concept categories accounting for (i.e., hidden causes generating) patterns in its possible observations. When a new animal is presented, the agent must first infer that the animal is novel and engage an unused “slot” in its state space (i.e., infer that a noisy, flat distribution better accounts for the new pattern of observations than any current state-observation mapping it knows), and then learn the new state-outcome contingencies over repeated observations. Thus, as we use the term, model expansion increases the number of hidden states the agent *uses*, but not the formal dimensions of the state space itself.

In these simulations, learning was implemented via updating (concentration) parameters for the model’s **A** matrix after each trial. For details of these free energy minimizing learning processes, please see (Friston et al., 2016a) as well as the left panel of Figure 2 and associated legend. An intuitive way to think about this belief updating process is that the strength of association between a concept and an observation is quantified simply by counting how often they are inferred to co-occur. This is exactly the same principle that underwrites Hebbian plasticity and long-term potentiation (Brown et al., 2010). Crucially, policies were restricted during learning, such that the agent could not select reporting actions; thus, learning was driven entirely by repeated exposure to different feature combinations. We evaluated successful learning in two ways. First, we compared the **A** matrix learned by the model to that of the generative process. Second, we disabled learning after various trial numbers (i.e., such that concentration parameters no longer accumulated) and enabled reporting. We then evaluated reporting accuracy with 20 trials for each of the 8 concepts.

As shown in Figure 4A, the **A** matrix (likelihood) mapping – learned by the agent – and the column for sturgeon in particular, strongly resembled that of the generative process. When first evaluating reporting, the model was 100 % accurate across 20 reporting trials, when exposed to a sturgeon (reporting accuracy when exposed to each of the other animals also remained at 100%) and first reached this level of accuracy after around 50 exposures to all 8 animals (with equal probability) (figure 4B). The agent also always chose to report specific categories (i.e., it never chose to only report bird or fish). Model performance was stable over 8 repeated simulations.

Crucially, during learning, the agent was not told which state was generating its observations. This meant that it had to solve both an inference and a learning problem. First, it had to infer whether a given feature combination was better explained by an existing concept, or by a concept that predicts features uniformly. In other words, it had to decide that the features were sufficiently different – from things it had seen before – to assign it a new hypothetical concept. Given that a novel state is only inferred when another state is not a better explanation, this precludes learning ‘duplicate’ states that generate the same patterns of observations. The second problem is simpler. Having inferred that these outcomes are caused by something new, the problem becomes one of learning a simple state-outcome mapping through accumulation of Dirichlet parameters.

To examine whether this result generalized, we repeated these simulations under conditions in which the agent had to learn more than one concept. When the model needed to learn one bird (parakeet) and one fish (minnow), the model was also able to learn the appropriate likelihood mapping for these 2 concepts (although note that, because the agent did not receive feedback about the state it was in during learning, the new feature mappings were often not assigned to the same columns as in the generative process; see figure 4A). Reporting also reached 100% accuracy, but required a notably greater number of trials. Across 8 repeated simulations, the mean accuracy reached by the model after 2000 trials was 98.75% (SD = 2%).

When the model needed to learn all 4 birds, performance varied somewhat more when the simulations were repeated. The learned likelihood mappings after 2000 trials always resembled that of the generative process, but with variable levels of precision; notably, the model again assigned different concepts to different columns relative to the generative process, as would be expected when the agent is not given feedback about the state it is in. Over 8 repeated simulations, the model performed well in 6 (92.50 % – 98.8 % accuracy) and failed to learn one concept in the other 2 (72.50 % accuracy in each) due to overgeneralization (e.g., mistaking parrot for Hawk in a majority of trials; i.e., the model simply learned that there are large birds). Figure 4A and 4B illustrate representative results when the model was successful (note: the agent never chose to report basic categories in the simulations where only 1 or 2 concepts needed to be learned).

To further assess concept learning, we also tested the agent’s ability to successfully avoid state duplication. That is, we wished to confirm that the model would only learn a new concept if actually presented with a new animal for which it did not already have a concept. To do so, we equipped the model with knowledge of seven out of the eight concept categories, and then repeatedly exposed it only to the animals it already knew over 80 trials. We subsequently exposed it to the eighth animal (Hawk) for which it did not already have knowledge over 20 additional trials. As can be seen in figure 5, the unused concept column was not engaged during the first 80 trials (bottom left and middle). However, in the final 20 trials, the agent correctly inferred that its current conceptual repertoire was unable to explain its new pattern of observations, leading the unused concept column to be adumbrated and the appropriate state-observation mapping to be learned (bottom right). We repeated these simulations under conditions in which the agent already had knowledge of six, five, or four concepts. In all cases, we observed that unused concept columns were never engaged inappropriately.

**Figure 5.**
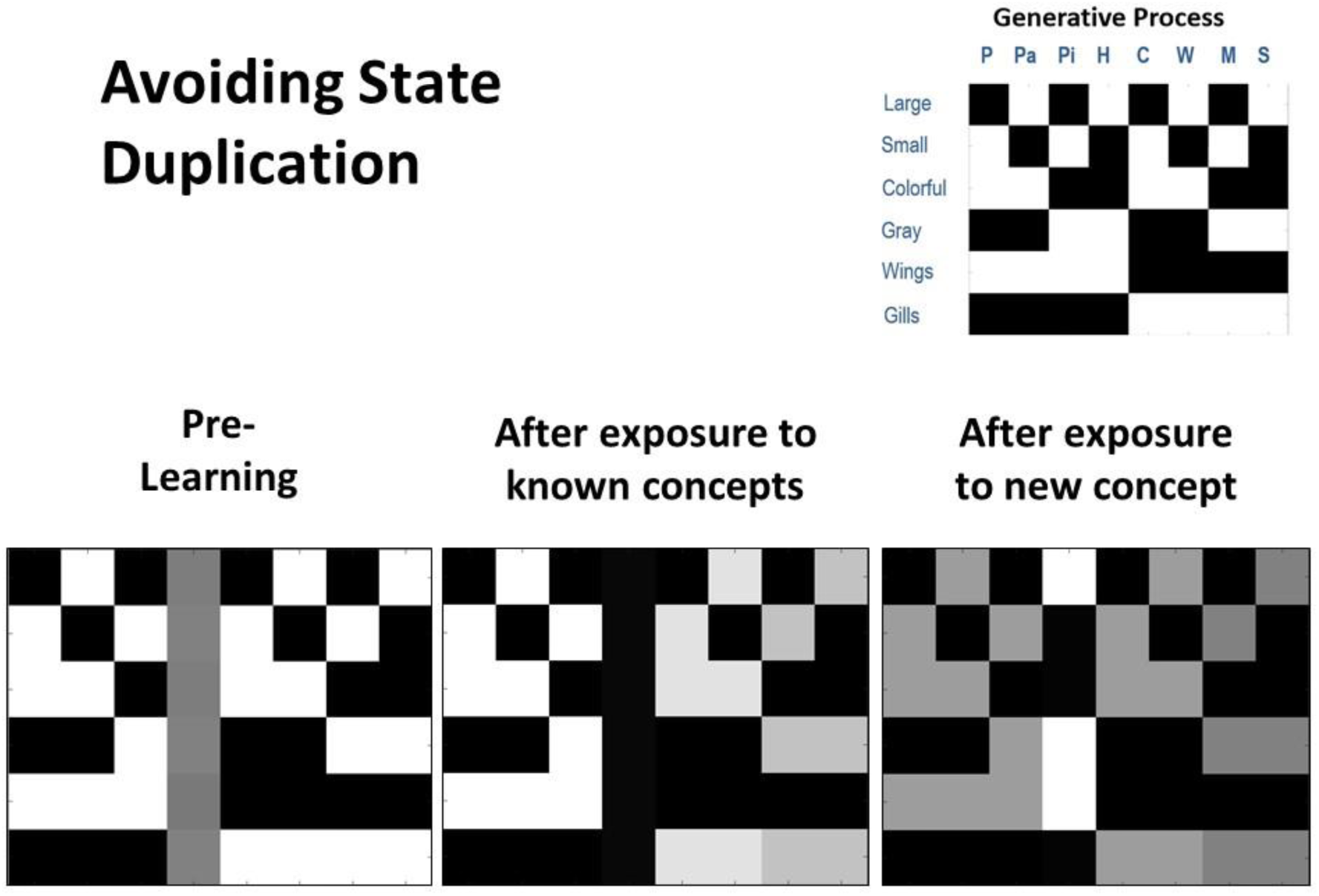
Illustration of representative simulation results when the agent had to avoid 548 inappropriately learning a new concept (i.e., avoid state duplication) after only being 549 exposed to animals for which it already had knowledge. Here the agent began with prior 550 knowledge about seven concept categories and was also equipped with an eighth column 551 that could be engaged to learn a new concept category (bottom left). The agent was then 552 presented with several instances of each of the seven animals that it already knew (80 trials 553 in total). In this simulation, the agent was successful in assigning each stimulus to an animal 554 concept it had already acquired and did not engage the unused concept ‘slot’ (bottom 555 middle). Finally, the agent was presented with a new animal (a hawk) that it did not already 556 know over 20 trials. In this case, the agent successfully engaged the additional column (i.e., 557 it inferred that none of the concepts it possessed could account for its new observations) 558 and learned the correct state-observation mapping (bottom right).

Crucially, these simulations suggest that adaptive concept learning needs to be informed by existing knowledge about other concepts, such that a novel concept should only be learned if observations cannot be explained with existing conceptual knowledge. Here, this is achieved via the interplay of inference and learning, such that agents initially have to infer whether to assign an observation to an existing concept, and only if this is not possible is an ‘open slot’ employed to learn about a novel concept.

#### Increasing granularity

Next, to explore the model’s ability to increase the granularity of its concept space, we first equipped the model with only the distinction between birds and fish (i.e., the rows of the likelihood mapping corresponding to color and size features were flattened in the same manner described above). We then performed the same procedure used in our previous simulations. As can be seen in Figure 6A (bottom left), the **A** matrix learned by the model now more strongly resembled that of the generative process. Figure 6B (bottom) also illustrates reporting accuracy and the proportion of basic and specific category reports as a function of trial number. As can be seen, the agent initially only reported general categories, and became sufficiently confident to report specific categories after roughly 50 – 100 trials, but its accuracy increased gradually over the next 1000 trials (i.e., the agent reported specific categories considerably before its accuracy improved). Across 8 repeated simulations, the final accuracy level reached was between 93% – 98% in 7 simulations, but the model failed to learn one concept in the 8th case, with 84.4% overall accuracy (i.e., a failure to distinguish between pigeon and parakeet, and therefore only learned a broader category of “small birds”).

**Figure 6.**
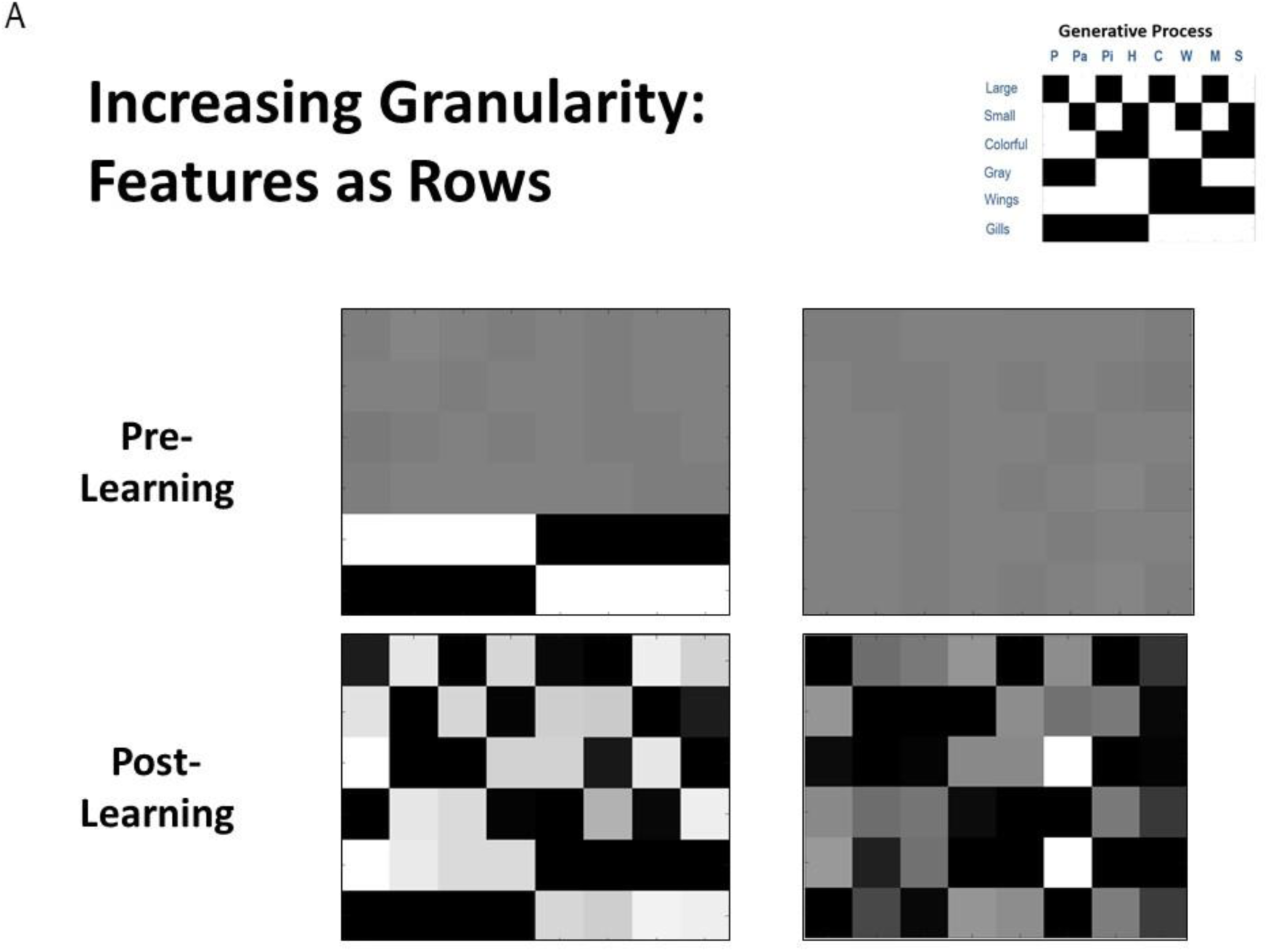
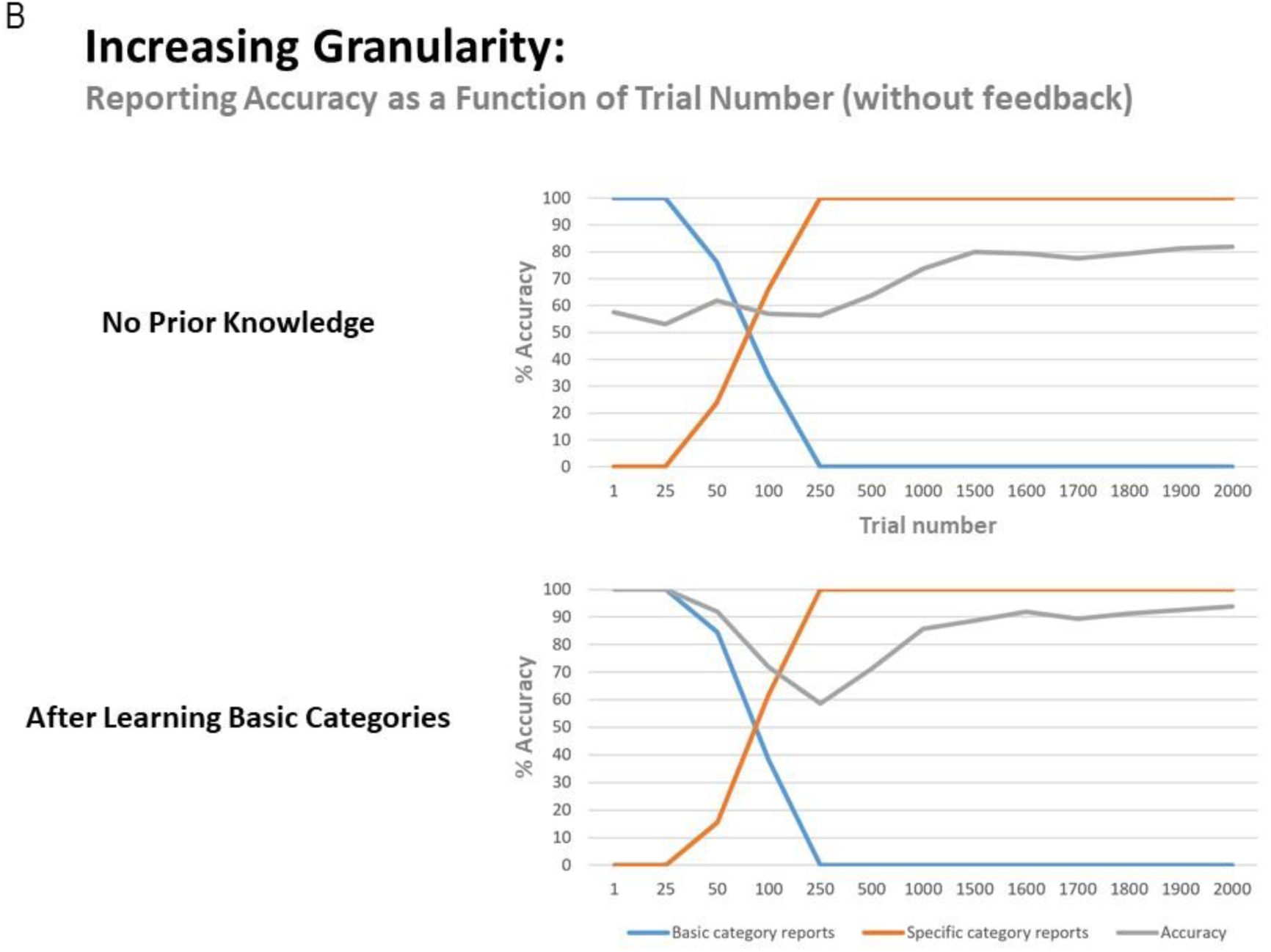
(A, left) Illustration of representative simulation results when the agent had to learn to increase the granularity of its concept space. Here the agent began with prior knowledge about the basic concept categories (i.e., it had learned the broad categories of “bird” and “fish”) but had not learned the feature patterns (i.e., rows) that differentiate different types of birds and fish. Post learning (i.e., after 2000 exposures), the agent did successfully learn all of the more granular concept categories, although again note that specific concepts were assigned to different columns then depicted in the generative process due to the unsupervised nature of the learning. (A, right) illustration of the analogous learning result when the agent had to learn all 8 specific categories without prior knowledge of the general categories. Although moderately successful, learning tended to be more difficult in this case. (B) Representative plots of reporting accuracy in each of the 2 learning conditions as a function of the number of exposures. As can be seen, when the model starts out with prior knowledge about basic categories, it slowly becomes sufficiently confident to start reporting the more specific categories, and its final accuracy is high. In contrast, while the agent that did not start out with any prior knowledge of the general categories also grew confident in reporting specific categories over time, its final accuracy levels tended to be lower. In both cases, the agent began reporting specific categories before it achieved significant accuracy levels, therefore showing some initial overconfidence.

To assess whether learning basic categories first was helpful in subsequently learning specific categories, we also repeated this simulation without any initial knowledge of the basic categories. As exemplified in figure 6A and 6B, the model tended to perform reasonably well, but most often learned a less precise likelihood mapping and reached a lower reporting accuracy percentage after 2000 learning trials (across 8 repeated simulations: mean = 81.21%, SD = 6.39%, range from 68.80% – 91.30%). Thus, learning basic concept categories first appeared to facilitate learning more specific concepts later.

Overall, these findings provide a proof of principle that this sort of active inference scheme can add concepts to a state space in an unsupervised manner (i.e., without feedback) based purely on (expected) free energy minimization. In this case, it was able to accomplish this starting from a completely uninformative likelihood distribution. It could also learn more granular concepts after already acquiring more basic concepts, and our results suggest that learning granular concepts may be facilitated by first learning basic concepts (e.g., as in currently common educational practices).

The novel feature of this generative model involved ‘building in’ a number of “reserve” hidden state levels. These initially had uninformative likelihood mappings; yet, if a new pattern of features was repeatedly observed, and the model could not account for this pattern with existing (informative) state-observation mappings, these additional hidden state levels could be engaged to improve the model’s explanatory power. This approach therefore accommodates a simple form of structure learning (i.e., model expansion).

### Integrating model expansion and reduction

We next investigated ways in which the form of effective model expansion simulated above could be combined with an analogous form of Bayesian model reduction (Friston et al., 2017b) – allowing the agent to adjust its model to accommodate new patterns of observations, while also precluding unnecessary conceptual complexity (i.e., over-fitting). To do so, we again allowed the agent to learn from 2000 exposures to different animals as described in the previous section – but also allowed the model to learn its ‘D’ matrix (i.e., accumulate concentration parameters reflecting prior expectations over initial states). This allowed an assessment of the agent’s probabilistic beliefs about which hidden state factor levels (animals) it had been exposed to. In different simulations, the agent was only exposed to some animals and not others. We then examined whether a subsequent model reduction step could recover the animal concepts presented during the simulation; eliminating those concepts that were unnecessary to explain the data at hand. The success of this 2-step procedure could then license the agent to “reset” the unnecessary hidden state columns after concept acquisition, which would have accrued unnecessary likelihood updates during learning. Doing so would allow the optimal ability for those “reserve” states to be appropriately engaged, if and when the agent was exposed to truly novel stimuli. Thus, as with our model expansion procedure, model reduction is not formally changing the dimensions of the agent’s state space. It instead prevents the unnecessary use of the agent’s “reserve” states so that they are only engaged when a novel animal is truly present.

The 2^nd^ step of this procedure was accomplished by applying Bayesian model reduction to the **D** matrix concentration parameters after learning. This is a form of post-hoc model optimization (Friston et al., 2016b, 2018) that rests upon estimation of a ‘full’ model, followed by analytic computation of the evidence that would have been afforded to alternative models (with alternative, ‘reduced’, priors) had they been used instead. Mathematically, this procedure is a generalization of things like automatic relevance determination (Friston et al., 2007; Wipf and Rao, 2007) or the use of the Savage Dickie ratio in model comparison (Cornish and Littenberg, 2007). It is based upon straightforward probability theory and, importantly, has a simple physiological interpretation; namely, synaptic decay and the elimination of unused synaptic connections. In this (biological) setting, the concentration parameters of the implicit Dirichlet distributions can be thought of as synaptic tags. For a technical description of Bayesian model reduction techniques and their proposed neural implementation, see (Hobson and Friston, 2012; Hobson et al., 2014; Friston et al., 2017b); see the left panel of Figure 2 for additional details).

The posterior concentration parameters were compared to the prior distribution for a full model (i.e., a flat distribution over 8 concepts) and prior distributions for possible reduced models (i.e., which retained different possible combinations of some but not all concepts; technically, reduced models were defined such that the to-be-eliminated concepts were less likely than the to-be-retained concepts). If Bayesian model reduction provided more evidence for one or more reduced models, the reduced model with the most evidence was selected.

Note: an alternative would be to perform model reduction on the **A** matrix (i.e., comparing the state-outcome mappings one has learned to alternative **A** matrices that reset specific columns to flat distributions), but this is more complex due to a larger space of possible reduced models.

In each simulation, we presented our agent with a different number of animals (between 7 and 2 animals), with 250 exposures to each animal in randomized order (mirroring the learning simulations shown in Figure 6). In Table 1, we present results of 100 repeated simulations using this setup and report the number of times that the winning model corresponded to each possible reduced model when compared to the full model. To provide a further sense of these results, Table 1 also reports the number of times the 2^nd^-best model instead corresponded to the number of causes in the generative process.

**Table 1.**
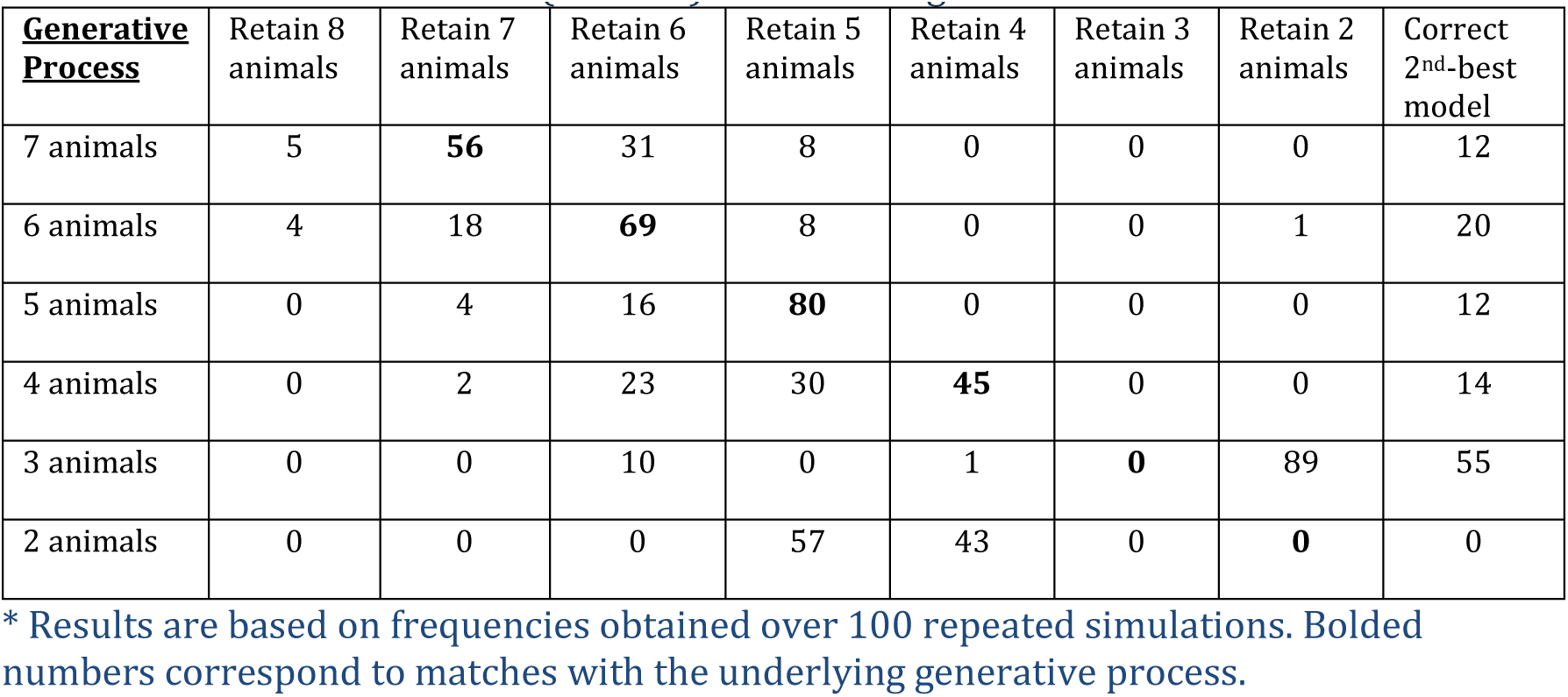
Frequency of winning models using Bayesian model reduction with different numbers of true hidden causes (animals) after learning.

| **Generative Process** | Retain 8 animals | Retain 7 animals | Retain 6 animals | Retain 5 animals | Retain 4 animals | Retain 3 animals | Retain 2 animals | Correct 2<sup>nd</sup>-best model |
| --- | --- | --- | --- | --- | --- | --- | --- | --- |
| 7 animals | 5 | **56** | 31 | 8 | 0 | 0 | 0 | 12 |
| 6 animals | 4 | 18 | **69** | 8 | 0 | 0 | 1 | 20 |
| 5 animals | 0 | 4 | 16 | **80** | 0 | 0 | 0 | 12 |
| 4 animals | 0 | 2 | 23 | 30 | **45** | 0 | 0 | 14 |
| 3 animals | 0 | 0 | 10 | 0 | 1 | **0** | 89 | 55 |
| 2 animals | 0 | 0 | 0 | 57 | 43 | 0 | **0** | 0 |

\* Results are based on frequencies obtained over 100 repeated simulations. Bolded numbers correspond to matches with the underlying generative process.

As can be seen there, the frequency with which model reduction successfully recovered the number of causes in the generative process varied depending on the number of different types of animals presented (Figure 7 illustrates examples of successful recovery in the case of generative processes with 7, 6, and 5 causes). In the presence of 4-7 animals, the most frequently winning model was the correct model. When the winning model was not the correct model, this was most often due to the agent learning simpler, coarser-grained representations (such as “large bird”, treating color as irrelevant; e.g., the 31/100 cases where a 6-animal model was retained after 7 different animals were presented). In some other cases, the agent instead retained overly fine-grained distinctions (such as the existence of distinct types of gray birds with different frequencies of being large vs. small; e.g., the several cases where a 5- or 6-animal model was retained when only 4 different animals were presented). In these cases where the incorrect model was favored, the correct model was often the model with the 2nd-most evidence. The difference in log evidence between the winning and 2nd-best model in such cases was also often small (average differences between -.40 and −1.23). In the approach outlined here, in the cases where model reduction was successful, this would correctly license the removal of changes in the model’s **A** and **D** matrix parameters for unobserved animals during learning in the previous trials (i.e., re-setting them to then again be available for future learning).

**Figure 7.**
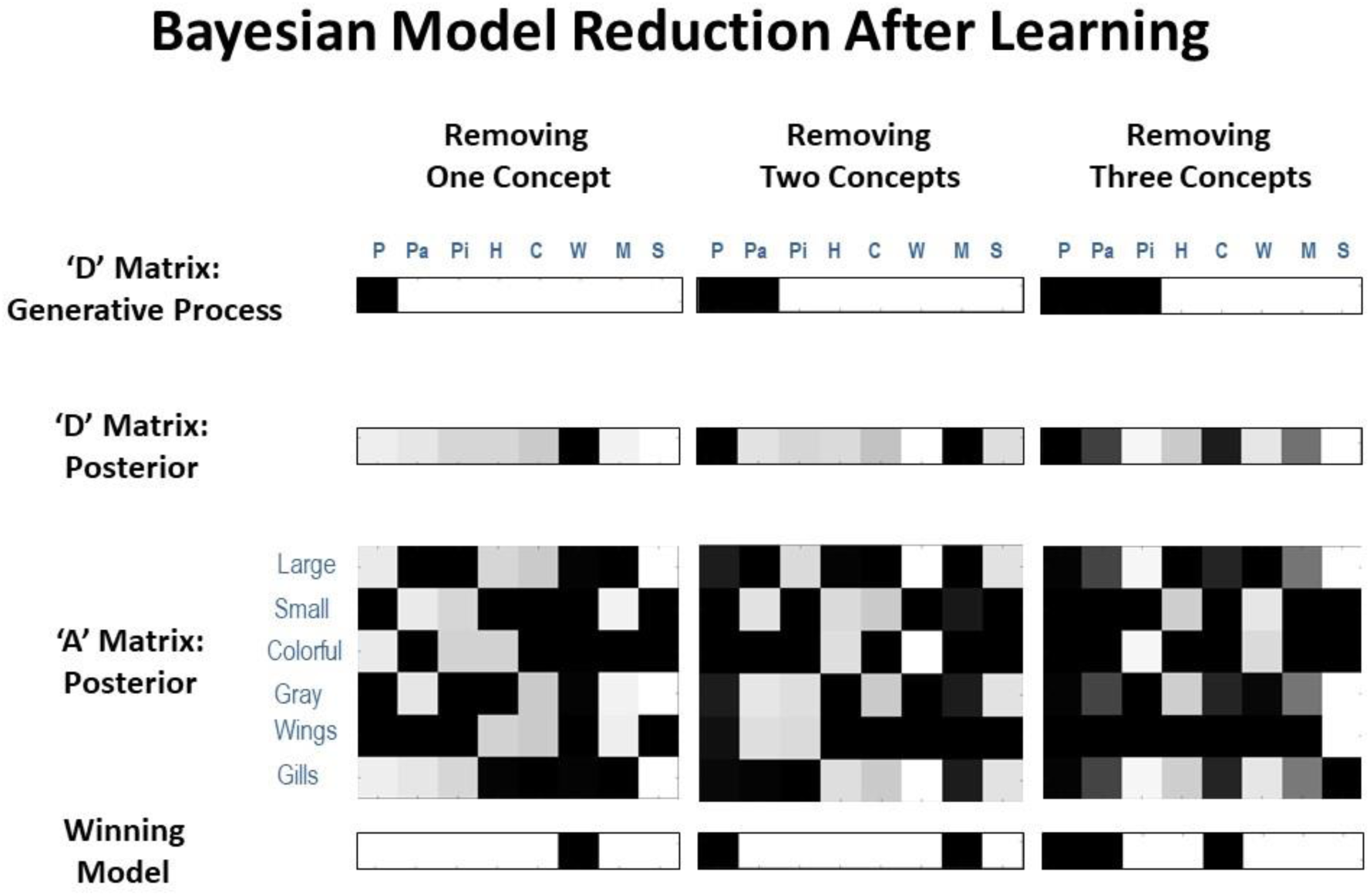
Representative illustrations of simulations in which the agent performed Bayesian model reduction after learning. In these simulations, the agent was first given 250 exposures per animal, where either 7, 6, or 5 animals were actually presented (i.e., illustrated in the top row, where only the white columns had nonzero probabilities in the generative process). In each case, model reduction was often successful at identifying the reduced model with the correct number of animal types presented (bottom row, where black columns should be removed) based on how much evidence it provided for the posterior distribution over hidden states learned by the agent (2^nd^ row). This would license the agent to reset the unneeded columns in its likelihood mapping (3^rd^ row) to their initial state (i.e., a flat distribution over features) such that they could be engaged if/when new types of animals began to be observed (i.e., as in the simulations illustrated in the previous sections).

When there were 3 causes to recover, the winning model most often only retained 2 animal representations (typically learning one more coarse-grained concept; e.g., “colorful fish”, ignoring size). However, the true 3-animal model had the 2^nd^-most evidence in the majority of unsuccessful cases (average log evidence difference with the winning model in these cases was −1.64).

When there were only 2 causes to recover (in this case, one bird and one fish), model reduction reliably failed to select the 2-animal model. Further inspection revealed that this was due to the agent learning – and failingly to subsequently remove – redundant state-outcome mappings across multiple (either 4 or 5) columns (i.e., it retained a fairly flat distribution over these states in **D**). Essentially, the agent learned 2 or 3 redundant mappings for ‘bird’ and 2 or 3 redundant mappings for ‘fish’. Poor performance in this case could be seen as unsurprising, as the presence of only 2 animals is quite different from the structural prior of 8 animals inherited by the agent prior to learning. Note, however, that despite a failure to retain only 2 concept columns, the resulting state-outcome mappings were coarse-grained in a manner similar to those of our simulated agent above that could only distinguish birds vs. fish (i.e., as in in the upper-left ‘pre-learning” matrix of Figure 6A, prior to granularity learning). This type of mapping only distinguishes 2 concepts (despite having redundant mappings across several columns) – and leads the agent to only report 2 coarse-grained categories (as shown in Figure 6B) – as would be expected if only familiar with 2 different animals distinguished by 1 feature. So, from another perspective, the agent successfully retained the type of mapping that reflects coarse-grained concept knowledge in our model and that allows for the appropriate coarse-grained reports.

When consider our overall model reduction results, it is worth noting that a model’s accuracy need not always correspond to its adaptiveness (McKay and Dennett, 2009; Al-Muhaideb and Menai, 2011; Gigerenzer and Gaissmaier, 2011; Baltieri and Buckley, 2019; Tschantz et al., 2019). In some cases, making either coarser- or finer-grained distinctions could be more adaptive for an organism depending on available resources and environmental/behavioral demands. It is also worth noting that, while we have used the terms ‘correct’ and ‘incorrect’ above to describe the model used to generate the data, we acknowledge that ‘all models are wrong’ (Box et al., 2005), and that the important question is not whether we can recover the ‘true’ process used to generate the data, but whether we can arrive at the simplest but accurate explanation for these data. The failures to recover the ‘true’ model highlighted above (especially in the cases where only coarser-grained representations were learned) may reflect that a process other than that used to generate the data could have been used to do so in a simpler way. Simpler here means we would have to diverge to a lesser degree from our prior beliefs in order to explain the data under a given model, relative to a more complex model. It is worth highlighting the importance of the word *prior* in the previous sentence. This means that the simplicity of the model is sensitive to our prior beliefs about it. To illustrate this, we repeated the same model comparisons as above, but with precise beliefs in an **A** matrix that complies with that used to generate the data. Specifically, we repeated the simulations above but only enabled **D** matrix learning (i.e., the model was already equipped with the **A** matrix of the generative process). In each case, Bayesian model reduction now uniquely identified the correct reduced model in 100% of repeated simulations.

Overall, these results demonstrate that – after a naïve model has effectively expanded its hidden state space to include likelihood mappings and initial state priors for a number of concept categories – Bayesian model reduction can subsequently be used with moderate success to eliminate any parameter updates accrued for redundant concept categories. In practice, the **A** and **D** concentration parameters for the redundant categories identified by model reduction could be reset to their default pre-learning values – and could then be re-engaged if new patterns of observations were repeatedly observed in the future. The variable performance of model reduction in our simulations appeared to be due to imperfect **A** matrix learning (recall that the analogous learning simulations in Figure 6 only led to 81% reporting accuracy on average). Another potential consideration is that, because **A** and **D** matrix learning occurred simultaneously, this may have also led to noisier accumulation of prior expectations over hidden states – as **D** matrix learning with a fully precise **A** matrix led to correct model reduction in every case tested (i.e., perhaps suggesting that this type of model reduction procedure could be improved by first allowing state-observation learning to proceed alone, then subsequently allowing the model to learn prior expectations over hidden states, which could then be used in model reduction).

### Can concept acquisition allow for generalization?

One important ability afforded by concept learning is generalization. In a final set of simulations, we asked if our model of concept knowledge could account for generalization. To do so, we altered the model such that it no longer reported what it saw, but instead had to answer a question that depended on generalization from particular cross-category feature combinations. Specifically, the model was shown particular animals and asked: “could this be seen from a distance?” The answer to this question depended on both size and color, such that the answer was yes only for colorful, large animals (i.e., assuming small or gray animals would blend in with the sky or water and be missed).

Crucially, this question was asked of animals that the model had not been exposed to, such that it had to generalize from knowledge it already possessed (see Figure 8). To simulate and test for this ability, we equipped the model’s **A** matrix with expert knowledge of 7 out of the 8 animals (i.e., as if these concepts had been learned previously, as in our simulations above). The 8^th^ animal was unknown to the agent, in that it’s likelihood mapping was set such that the 8^th^ animal state “slot” predicted all observations equally (i.e., with a small amount of Gaussian noise, as above). In one variant, the model possessed all concepts except for “parrot,” and it knew that the answer to the question was yes for “whale shark” but not for any other concept it knew. To simulate one-shot generalization, learning was disabled and a parrot (which it had never seen before) was presented 20 times to see if it would correctly generalize and answer “yes” in a reliable manner. In another variant, the model had learned all concepts except “minnow” and was tested the same way to see if it would reliably provide the correct “no” response.

**Figure 8.**
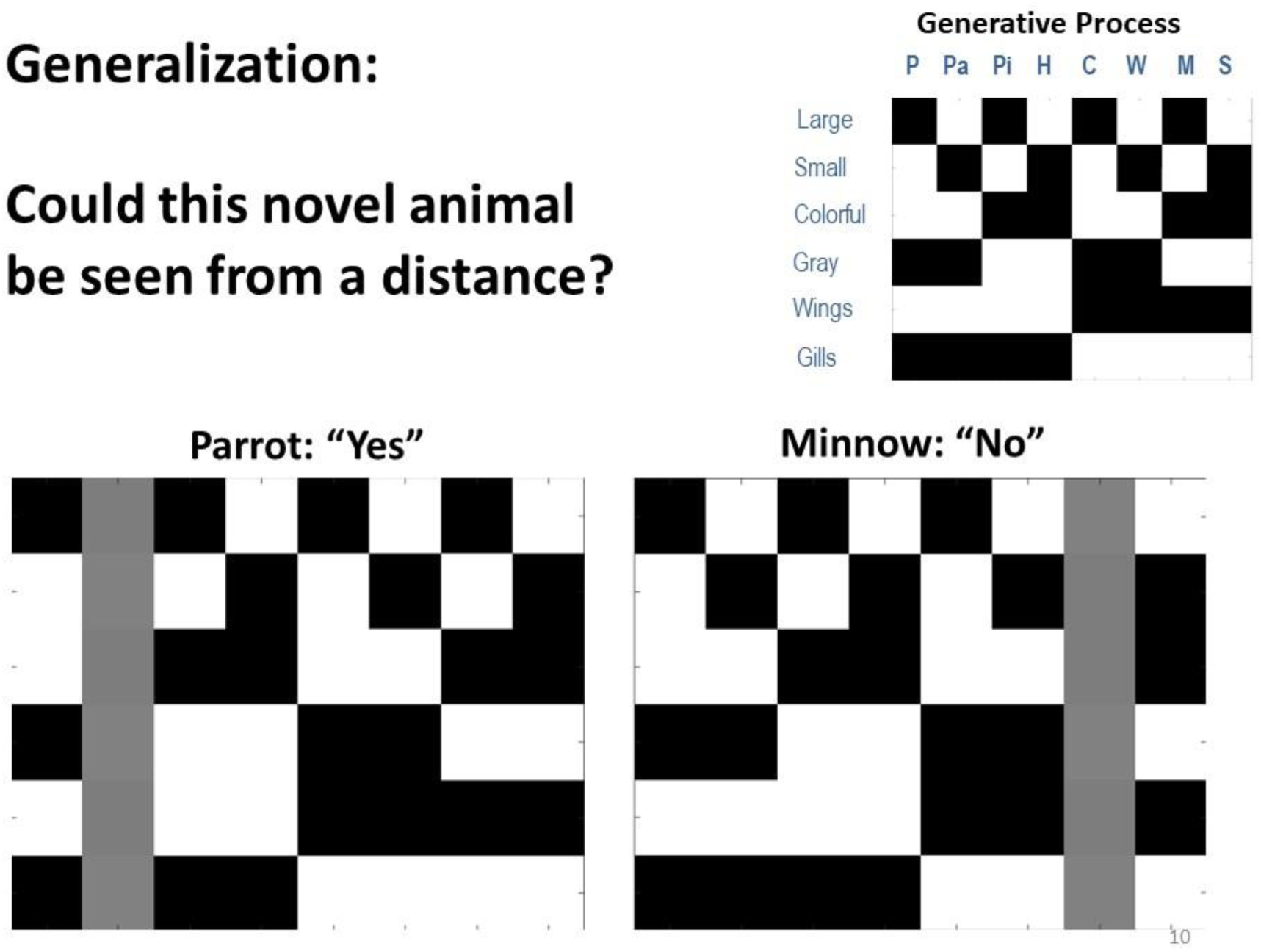
Depiction of simulations in which we tested the agent’s ability to generalize from prior knowledge and correctly answered questions about new animals to which it had not previously been exposed. In the simulations, the generative model was modified so that the agent instead chose to report either “yes” or “no” to the question: “could this animal be seen from a distance?” Here, the answer was only yes if the animal was both large and colorful. We observed that when the agent started out with no knowledge of parrots it still correctly answered this question 100% of the time, based only on its knowledge of other animals. Similarly, when it started with no knowledge of minnows, it also correctly reported “no” 100% of the time. Thus, the agent was able to generalize from prior knowledge with no additional learning.

Here, we observed that in both of these cases (as well as all others we tested) the model generalized remarkably well. It answered “yes” and “no” correctly in 100% of trials. Thus, the agent did not simply possess concepts to explain things it saw. It instead demonstrated generalizable knowledge and could correctly answer questions when seeing a novel stimulus.

### Open questions and relation to other theoretical accounts of concept learning

As our simulations show, this model allows for learning novel concepts (i.e., novel hidden states) based on assigning one or more ‘open slots’ that can be utilised to learn novel feature combinations. In a simple example, we have shown that this setup offers a potential computational mechanism for ‘model expansion’; i.e., the process of effectively expanding a state space to account for novel instances in perceptual categorisation. We also illustrated how this framework can be combined with model reduction, which may be a mechanism for ‘re-setting’ these open slots based on recent experience.

This provides a first step towards understanding how agents flexibly expand or reduce their model to adapt to ongoing experience. Yet, several open questions remain, and for the approach we illustrate to address real-world problems in future active inference research, these issues will need to be examined and integrated with insights from previous work that has already grappled with them. For example, the proposed framework resonates with previous similarity-based accounts of concept learning. Previous work has proposed a computational framework for arbitrating between assigning an observation to a previously formed memory or forming a novel (hidden) state representation (Gershman et al., 2017), based on evidence that this observation was sampled from an existing or novel latent state. This process is conceptually similar to our application of Bayesian model reduction over states. In the present framework, concept learning relies on a process based on inference and learning. First, agents have to *infer* whether ongoing observations can be sufficiently explained by existing conceptual knowledge – or speak to the presence of a novel concept that motivates the use of an ‘open slot’. This process is cast as inference on (hidden) states. Second, if the agent infers that there is a novel concept that explains current observations, it has to *learn* about the specific feature configuration of that concept (i.e., novel state). This process highlights the interplay between inference, which allows for the acquisition of knowledge on a relatively short timescale, and learning, which allows for knowledge acquisition on a longer and more stable timescale.

Similar considerations apply to the degree of ‘similarity’ of observations. In the framework proposed here, we have assumed that the feature space of observations is already learned and fixed. However, these feature spaces have to be learned in the first place, which implies learning the underlying components or feature dimensions that define observations. This relates closely to notions of structure learning as dimensionality reduction based on covariance between observations, as prominently discussed in the context of spatial navigation (Dordek et al., 2016; Stachenfeld et al., 2016; Behrens et al., 2018; Whittington et al., 2018).

Another important issue is how such abstract conceptual knowledge is formed across different contexts or tasks. For example, the abstract concept of a ‘bird’ will be useful for learning about the fauna in a novel environment, but specific types of birds – tied to a previous context – might be less useful in this regard. This speaks to the formation of abstract, task-general knowledge that results from training across different tasks, as recently discussed in the context of meta-reinforcement learning (Wang et al., 2016; Ritter et al., 2018) with a putative link to the prefrontal cortex (Wang et al., 2018). In the present framework, such task-general knowledge would speak to the formation of a hierarchical organisation that allows for the formation of conceptual knowledge both within and across contexts. Also note that our proposed framework depends on a pre-defined state space, including a pre-defined set of ‘open slots’ that allow for novel context learning. The contribution of the present framework is to show how these ‘open slots’ can be used for novel concept learning and be re-set based on model reduction. It will be important to extend this approach towards learning the structure of these models in the first place, including the appropriate number of ‘open slots’ (i.e., columns of the A-matrix) for learning in a particular content domain and the relevant feature dimensions of observations (i.e., rows of A-matrix). (Note: In addition to ontogenetic learning, in some cases structural priors regarding the appropriate number of open slots [and relevant feature inputs for learning a given state space of open slots] might also reflect inherited [i.e., genetically/developmentally pre-specified] patterns of structural neuronal connectivity – based on what was adaptive within the evolutionary niche of a given species – which could then be modified based on subsequent experience.)

This corresponds to a potentially powerful and simple application of Bayesian model reduction, in which candidate models (i.e., reduced forms of a full model) are readily identifiable based upon the similarity between the likelihoods conditioned upon different hidden states. If two or more likelihoods are sufficiently similar, the hidden states can be merged (by assigning the concentration parameters accumulated during experience-dependent learning to one or other of the hidden states). The ensuing change in model evidence scores the reduction in complexity. If this reduction is greater than the loss of accuracy – in relation to observations previously encountered – Bayesian model reduction will, effectively, merge one state into another; thereby freeing up a state for the learning of new concepts. We will demonstrate this form of structure learning via Bayesian model reduction in future work.

We must also highlight here that cognitive science research on concept and category learning has a rich empirical and theoretical history, including many previously proposed formal models. While our primary focus has been on using concept learning as an example of a more general approach by which state space expansion and reduction can be implemented within future active inference research, it is important to recognize this previous work and highlight where it overlaps with the simulations we’ve presented. For example, our results suggesting that first learning general categories facilitates the learning of more specific categories relates to both classic and contemporary findings showing that children more easily acquire “basic” and “superordinate” (e.g., dog, animal) concepts before learning “subordinate” (e.g., chihuahua) concepts (Mervis and Rosch, 1981; Murphy, 2016), and that this may involve a type of “bootstrapping” process (Beck, 2017).

Complementary work has also highlighted ways in which learning new words during development can invoke a type of “placeholder” structure, which then facilitates the acquisition of a novel concept – which bears some resemblance to our notion of blank “concept slots” that can subsequently acquire meaningful semantics (Gelman and Roberts, 2017).

There is also a series of previously proposed formalisms within the literature on category learning. For example, two previously proposed models – the “rational model” (Anderson, 1991; Sanborn et al., 2010) and the SUSTAIN model (Love et al., 2004) – both describe concept acquisition as involving cluster creation mechanisms that depend on statistical regularities during learning and that use probabilistic updating. The updating mechanisms within SUSTAIN are based on surprise/prediction-error in the context of both supervised and unsupervised learning. This model also down-weights previously created clusters when their associated regularities cease to be observed in recent experience. Although not built in intentionally, this type of mechanism emerges naturally within our model in two ways. First, when a particular hidden state ceases to be inferred, concentration parameters will accumulate to higher values for other hidden states in the **D** matrix, reflecting relatively stronger prior expectations for hidden states that continue to be inferred – which would favor future inference of those states over those absent from recent experience. Second, if one pattern of observations were absent from recent experience (while other patterns continued to be observed), concentration parameters in the **A** matrix would also accumulate to higher values for patterns that continued to be observed – resulting in relatively less confidence in the state-outcome mapping for the less-observed pattern. (However, with respect to this latter mechanism, so long as this mapping was sufficiently precise and distinct from others [i.e., it had previously been observed many times farther in the past], this would not be expected to prevent successful inference if this pattern were observed again.)

It is also worth highlighting that, as our model is intended primarily as a proof of concept and a demonstration of an available model expansion/reduction approach that can be used within active inference research, it does not explicitly incorporate some aspects – such as top-down attention – that are of clear importance to cognitive learning processes, and that have been implemented in previous models. For example, the adaptive resonance theory (ART) model (Grossberg, 1987) was designed to incorporate top-down attentional mechanisms and feedback mechanisms to address a fundamental knowledge acquisition problem – the temporal instability of previously learned information that can occur when a system also remains sufficiently plastic to learn new (and potentially overlapping) information. While our simulations do not explicitly incorporate these additional complexities, there are clear analogues to the top-down and bottom-up feedback exchange in ART within our model (e.g., the prediction and prediction-error signaling within the neural process theory associated with active inference). ART addresses the temporal instability problem primarily through mechanisms that learn top-down expectancies that guide attention and match them with bottom-up input patterns – which is quite similar to the prior expectations and likelihood mappings used within active inference.

As an emergent property of the “first principles” approach in active inference, our model therefore naturally incorporates the top-down effects in ART simulations, which have been used to account for known context effects on categorical perception within empirical studies (McClelland and Rumelhart, 1981). This is also consistent with more recent work on cross-categorization (Shafto et al., 2011), which has shown that human category learning is poorly accounted for by both a purely bottom-up process (attempting to explain observed features) and a purely top-down approach (involving attention-based feature selection) – and has instead used simulations to show that a Bayesian joint inference model better fits empirical data.

Other proposed Bayesian models of concept learning have also had considerable success in predicting human generalization judgments (Goodman et al., 2008). The proof of concept model presented here has not been constructed to explicitly compete with such models. It will be an important direction for future work to explore the model’s ability to scale up to handle more complex concept learning problems. Here we simply highlight that the broadly Bayesian approach within our model is shared with other models that have met with considerable success – supporting the general plausibility of using this approach within active inference research to model and predict the neural basis of these processes (see below).

### Potential advantages of the approach

The present approach may offer some potential theoretical and empirical advantages in comparison to previous work. One theoretical advantage corresponds to the parsimony of casting this type of structure learning as an instance of Bayesian model selection. When integrated with other aspects of the active inference framework, this means that perceptual inference, active learning, and structure learning are all expressions of the same principle; namely, the minimization of variational free energy, over three distinct timescales. Structure learning is particularly interesting from this perspective, as it occurs both during an individual’s lifetime, and over the longer timescale of natural selection; which implements a form of structure learning by selecting among phenotypes that entail alternative models. It also highlights the importance of these nested timescales within an individual’s lifetime, in that active learning must proceed through repeated perception of the consequences of action, and structure learning must proceed by 1) accumulating evidence that the state-outcome mappings one has already learned are insufficient to account for new observations (entailing the need for model expansion), and 2) using model reduction to remove actively learned state-outcome mappings that are unnecessary to account for past observations. A second, related theoretical advantage is that, when this type of structure learning is cast as Bayesian model selection/reduction, there is no need to invoke additional procedures or schemes (e.g., nonparametric Bayes or ‘stick breaking’ processes; (Gershman and Blei, 2012)). Instead, a generative model with the capacity to represent a sufficiently complex world will automatically learn causal structure in a way that contextualizes active inference within active learning, and active learning within structure learning.

Based on the process theories summarized in Figure 2, the present model would predict that the brain contains “reserve” cortical columns and synapses (most likely within secondary sensory and association cortices) available to capture new patterns in observed features. To our knowledge, no direct evidence supporting the presence of unused cortical columns in the brain has been observed, although the generation of new neurons (with new synaptic connections) is known to occur in the hippocampus (Chancey et al., 2013). “Silent synapses” have also been observed in the brain, which does appear consistent with this prediction; such synapses can persist into adulthood and only become activated when new learning becomes necessary (e.g., see (Kerchner and Nicoll, 2008; Chancey et al., 2013; Funahashi et al., 2013)). One way in which this idea of “spare capacity” or “reserve” cortical columns might be tested in the context of neuroimaging would be to examine whether greater levels of neural activation – within conceptual processing regions – are observed after learning additional concepts, which would imply that additional populations of neurons become capable of being activated. In principle, single-cell recording methods might also test for the presence of neurons that remain at baseline firing rates during task conditions, but then become sensitive to new stimuli within the relevant conceptual domain after learning.

Figure 9 provides a concrete example of two specific empirical predictions that follow from simulating the neural responses that should be observed within our concept learning task under these process theories. In the left panel, we plot the firing rates (darker = higher firing rate) and local field potentials (rate of change in firing rates) associated with neural populations encoding the probability of the presence of different animals that would be expected across a number of learning trials. In this particular example, the agent began with knowledge of the basic categories of ‘bird’ and ‘fish,’ but needed to learn the eight more specific animal categories over 50 interleaved exposures to each animal (only 10 equally spaced learning trials involving the presentation of a parakeet are shown for simplicity). As can be seen, early in learning the firing rates and local field potentials remain at baseline levels; in contrast, as learning progresses, these neural responses take a characteristic shape with more and more positive changes in firing rate in the populations representing the most probable animal, while other populations drop further and further below baseline firing rates.

**Figure 9.**
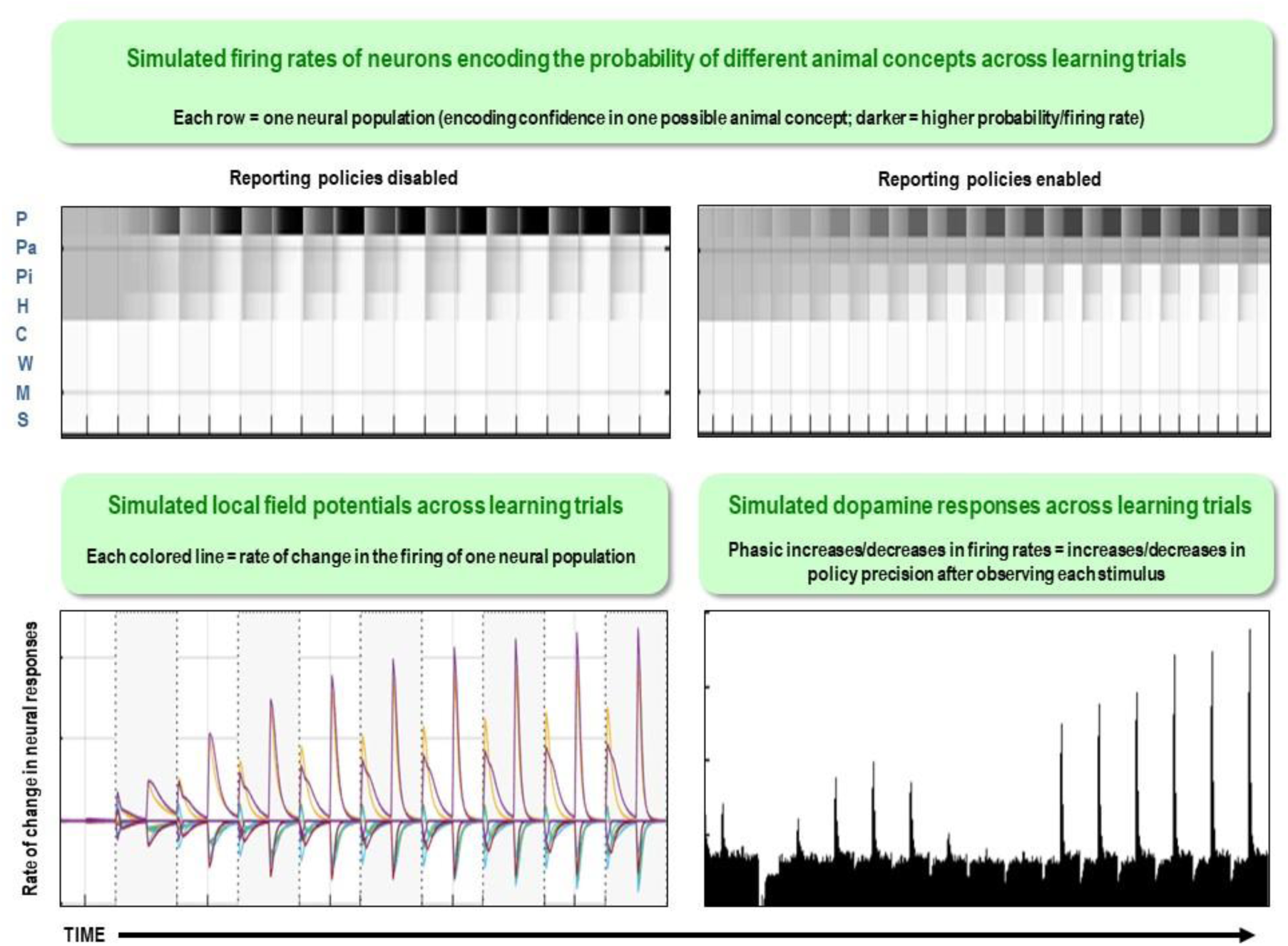
Simulated neuronal firing rates, local field potentials, and dopaminergic responses across learning trials based on the neural process theory associated with active inference that is summarized in Figure 2. The top left panel displays the predicted firing rates (darker = higher firing rate) of neural populations encoding the probability of each hidden state over 50 interleaved exposures to each animal (only 10 equally spaced learning trials involving the presentation of a parakeet are shown for simplicity) in the case where the agent starts out with knowledge of the basic animal categories but must learn the more specific categories. As can be seen, initially each of the four neural populations encoding possible bird categories (i.e., one row per possible category) have equally low firing rates (gray); as learning continues, firing rates increase for the ‘parakeet’ population and decrease for the others. The bottom left panel illustrates the predicted local field potentials (based on the rate of change in firing rates) that would be measured across the task. The top right panel displays the predicted firing rates of neural populations in an analogous simulation in which reporting policies were enabled (for clarity of illustration, we here show 12 equally spaced learning trials for parakeet over 120 total trials). Enabling policy selection allowed us to simulate the phasic dopaminergic responses (reporting changes in the precision of the probability distribution over policies) predicted to occur across learning trials; here the agent first becomes confident in its ability to correctly report the general animal category upon observing a stimulus, then becomes unsure about reporting specific versus general categories, and then becomes confident in its ability to report the specific categories.

The right panel depicts a similar simulation, but where the agent was allowed to self-report what it saw on each trial (for clarity of illustration, we here show 12 equally spaced learning trials for parakeet over 120 total trials). Enabling policy selection allowed us to simulate expected phasic dopamine responses during the task, corresponding to changes in the precision of the probability distribution over policies after observing a stimulus on each trial. As can be seen, during early trials the model predicts small firing rate increases when the agent is confident in its ability to correctly report the more general animal category after observing a new stimulus, and firing rate decreases when the agent becomes less confident in one policy over others (i.e., as confidence in reporting the specific versus general categories becomes more similar). Larger and larger phasic dopaminergic responses are then expected as the agent becomes more and more confident in its ability to correctly report the specific animal category upon observing a new stimulus. It will be important for future neuroimaging studies to test these predictions in this type of concept learning/stimulus categorization task.

## Discussion

The Active Inference formulation of concept learning presented here demonstrates a simple way in which a generative model can effectively expand its state-space to acquire both basic and highly granular knowledge of the hidden states/causes in its environment. In comparison to previous theoretical work using active inference (e.g., (Schwartenbeck et al., 2015; Mirza et al., 2016; Parr and Friston, 2017)), the novel aspect of our model was that it was further equipped with “reserve” hidden states initially devoid of content (i.e., these states started out with uninformative likelihood mappings that predicted all outcomes with roughly equal probability). Over multiple exposures to different stimuli, these hidden states came to acquire conceptual content that captured distinct statistical patterns in the features of those stimuli. This was accomplished via the model’s ability to infer when its currently learned hidden states were unable to account for a new observation, leading an unused hidden state column to be engaged that could acquire a new state-observation mapping. This provides a simple illustration of how this approach could be used in active inference research and applied to more complex structure learning problems.

Crucially, the model was able to start with some concepts and then expand its representational repertoire to learn others – but would only do so when a novel stimulus was observed and detected (i.e., when the model inferred that the flat/noisy state-outcome mapping of an unused hidden state column better accounted for its current observations than the existing concept mappings it had already acquired – a type of “novel concept detection”). This is conceptually similar to nonparametric Bayesian learning models, such as the “Chinese Room” process and the “Indian Buffet” process, that can also infer the need to invoke additional hidden causes with additional data (Gershman and Blei, 2012). These statistical learning models do not need to build in additional “category slots” for learning as in our model and can, in principle, entertain infinite state spaces. On the other hand, it is less clear at present how the brain could implement this type of learning. An advantage of our model is that learning depends solely on biologically plausible Hebbian mechanisms (for a possible neural implementation of model reduction, see (Hobson and Friston, 2012; Hobson et al., 2014; Friston et al., 2017b)).

The distinction between nonparametric Bayesian learning and the current active learning scheme may be important from a neurodevelopmental perspective as well. In brief, structure learning in this paper starts with a generative model with a type of structural prior reflecting a specific amount of built in ‘spare capacity’, where uncommitted or naive conceptual ‘slots’ are used to explain the sensorium, during optimization of free energy or model evidence. In contrast, nonparametric Bayesian approaches add new slots when appropriate. One might imagine that neonates are equipped with brains with ‘spare capacity’ (Baker and Tenenbaum, 2014) that is progressively leveraged during neurodevelopment, much in the spirit of curriculum learning (Al-Muhaideb and Menai, 2011). This suggestion appears consistent with previous work demonstrating varying levels of category learning ability across the lifespan, which has previously been formally modeled as an individual difference in values of a parameter constraining the ability to form new clusters in response to surprising events (Love and Gureckis, 2007) – which bears similarity to the idea of capacity limitations arising from finite numbers of concept slots in our model.

In this sense, the current approach to structure learning, and its implementation for effectively expanding a state space (i.e., increasing the number hidden states an agent actively uses to understand its environment) can be understood as a specific type of active learning with generative models that are equipped with a large number of available hidden states capable of acquiring content, which are then judiciously reduced/reset – via a process of Bayesian model reduction. Furthermore, as in the acquisition of expertise, our model can also begin with broad category knowledge and then subsequently learn finer-grained within-category distinctions, which has received less attention from the perspective of the aforementioned models. Reporting broad versus specific category recognition is also a distinct aspect of our model – driven by differing levels of uncertainty and an expectation (preference) not to incorrectly report a more specific category.

Our simulation results also demonstrated that, when combined with Bayesian model reduction, the model can guard against learning too many categories during model expansion – often retaining only the number of hidden causes actually present in its environment – and thus keep “reserve” hidden states available for learning about new causes if or when they appear. With perfect “expert” knowledge of the possible animal types it could observe (i.e., fully precise likelihood mappings matching the generative process) this was true in general. With an imperfectly learned likelihood mapping, model reduction performed moderately well when only a few possible concepts needed to be removed, but performance worsened when larger numbers of concepts needed to be removed. This is perhaps unsurprising, as one might expect worse performance when the true number of hidden causes in the environment diverges more strongly from the agent’s structural priors (e.g., when 5 vs. 2 animals are presented to an agent with a structural prior for up to 8 animals). It would be interesting to test whether a similar pattern of learning is present in humans.

Neurobiological theories associated with Active Inference also make predictions about the neural basis of this process (Hobson and Friston, 2012; Hobson et al., 2014). Specifically, during periods of rest (e.g., daydreaming) or sleep, it is suggested that, because sensory information is down-weighted, learning is driven mainly by internal model simulations (e.g., as appears to happen in the phenomenon of hippocampal replay; (Pfeiffer and Foster, 2013; Feld and Born, 2017; Lewis et al., 2018)); this type of learning can accomplish a model reduction process in which redundant model parameters are identified and removed to prevent model over-fitting and promote selection of the most parsimonious model that can successfully account for previous observations. This is consistent with work suggesting that, during sleep, many (but not all) synaptic strength increases acquired in the previous day are attenuated (Tononi and Cirelli, 2014). The role of sleep and daydreaming in keeping “reserve” representational resources available for model expansion could therefore be especially important to concept learning – consistent with the known role of sleep in learning and memory (Stickgold et al., 2001; Walker and Stickgold, 2010; Perogamvros and Schwartz, 2012; Ackermann and Rasch, 2014; Feld and Born, 2017).

It is worth highlighting that, because spare capacity will have finite limits, it follows that state space expansion will also have limits (although these limits could be extremely high in complex biological systems). To manage such limitations effectively, an agent would need to come equipped with adaptive structural priors about the maximum number of possible categories (hidden causes) that could be ‘out there’ to learn about in a given domain (e.g., at each relevant timescale, level of abstraction, level of multimodal integration, etc.). Such priors for state space size – in a given domain – could influence learning in important ways. Most straightforwardly, having priors for a state space with too few possible causes would prevent adaptive (granular) learning. In contrast, it is less obvious what the effects of priors for a state space with too many possible causes would be. In the simulations above, our model could avoid state duplication – a case where, even when there was a prior for 8 possible categories, the model did not inappropriately learn an 8^th^ concept when only exposed to known hidden causes. This is because the agent only infers a new cause – and starts to learn a new state-outcome mapping – when a flat (noisy) mapping provides a better explanation for observations than previously learned mappings. This would therefore not be expected to occur in the presence of a familiar hidden cause, even with priors for a much larger state space (i.e., a greater number of empty ‘slots’). However, in our simulations of model reduction – where the agent started out with no prior category knowledge and had to learn the correct number of causes – the model was less often successful when fewer causes were present. In some cases, this was because the additional empty ‘slots’ allowed the agent to learn overly fine-grained or redundant mappings that model reduction failed to remove. This demonstrates one way in which having prior expectations that there are too many causes to learn about could also have detrimental effects during early development – perhaps suggesting the need for mechanisms that initially restrict spare capacity during early learning.

Another emergent feature of our model was its ability to generalize prior knowledge to new stimuli to which it had not previously been exposed. In fact, the model could correctly generalize upon a single exposure to a new stimulus – a type of “one-shot learning” capacity qualitatively similar to that observed in humans (Landau et al., 1988; Markman, 1989; Xu and Tenenbaum, 2007b). While it should be kept in mind that the example we have provided is very simple, it demonstrates the potential usefulness of this novel approach. Some other prominent approaches in machine-learning (e.g., deep learning) tend to require larger amounts of data (Geman et al., 1992; Lecun et al., 1998; Hinton et al., 2012; LeCun et al., 2015; Mnih et al., 2015), and do not learn the rich structure that allows humans to use concept knowledge in a wide variety of generalizable functions (Osherson and Smith, 1981; Barsalou, 1983; Biederman, 1987; Ward, 1994; Feldman, 1997; Markman and Makin, 1998; Williams and Lombrozo, 2010; Jern and Kemp, 2013). Other recent hierarchical Bayesian approaches in cognitive science have made progress in this domain, however, by modeling concepts as types of probabilistic programs using Bayesian program learning (Ghahramani, 2015; Goodman et al., 2015; Lake et al., 2015), and by using models that integrate hierarchical Bayesian approaches with deep learning (Salakhutdinov et al., 2013a) – both allowing learning from a smaller number of training examples. Unlike many of the other approaches mentioned above, these models are not based on clustering algorithms and represent an important step forward within the context of Bayesian models of cognition.

It is important to note that this model is deliberately simple and is meant only to represent a proof of principle that categorical inference and conceptual knowledge acquisition can be modeled within this particular neurocomputational framework, and to present this approach as a potentially useful tool in future active inference research. We chose a particular set of feature combinations to illustrate this, but it remains to be demonstrated that learning in this model would be equally successful with a larger feature space and set of learnable hidden causes. Due to limited scope, we have also not thoroughly addressed all other overlapping lines of research. For example, work on exemplar models of concepts has also led to other computational approaches. As one example, the EBRW model (Nosofsky and Palmeri, 1997) has demonstrated ways of linking exemplar learning to drift diffusion models. Another model within this line of research is the ALCOVE model (Nosofsky et al., 1994) – an error-driven connectionist model of exemplar-based category learning that employs selective attention and learns attentional weights (this model also built on earlier work; see (Nosofsky, 2011)). Yet another connectionist model with some conceptual overlap to our own is the DIVA model, which learns categories by recoding observations as task-constrained principle components and uses model fit for subsequent recognition (Kurtz, 2007). It will be important in future work to examine the strengths and limitations of a scaled-up version of our approach in relation to these models.

Yet another topic for future work would be the expansion of this type of model to context-specific learning (e.g., with an additional hidden state factor for encoding distinct contexts). In such cases, regularities in co-occurring features differ in different contexts and other cues to context may not be directly observable (e.g., the same species of bird could be a slightly different color or size in different parts of the world that otherwise appear similar) – creating difficulties in inferring when to update previously learned associations and when to instead acquire competing associations assigned to new contexts. At present, it is not clear whether the approach we have illustrated would be successful at performing this additional function, although the process of inferring the presence of a new hidden state level in a second hidden state factor encoding context would be similar to what we have shown within a single state factor (for related work on context-dependent contingency learning, see (Gershman et al., 2013, 2017)). Another point worth highlighting is that we have made particular choices with regard to various model parameters and the number of observations provided during learning. Further investigations of the space of these possible parameter settings will be important.

With this in mind, however, our current modelling results could offer additional benefits. For example, the model’s simplicity could be amenable to empirical studies of saccadic eye movements toward specific features during novel category learning (e.g. following the approach of (Mirza et al., 2018)). This approach could also be combined with measures of neural activity in humans or other animals, allowing more direct tests of the neural predictions highlighted above. In addition, the introduction of exploratory, novelty-seeking, actions could be used to reduce the number of samples required for learning, with agents selecting those data that are most relevant.

Introducing actions allowing selective data sampling also brings with it other interesting opportunities for using the general modeling approach used above. In our model, the agent learned passively through observations (as a practical means of preventing learning through feedback). Yet, while infants may initially be somewhat restricted to passively observing their environment, observations quickly come under the control of action (perhaps bootstrapping in a useful way on passive observations to first acquire sensorimotor contingencies; (Deci and Ryan, 1985; Oudeyer, 2007; Schmidhuber, 2010; Barto et al., 2013; Baltieri and Buckley, 2019)) – opening the possibility for both beneficial and detrimental effects on learning. By incorporating action into our active inference model, further questions could be addressed. For example, under what circumstances does choosing where to allocate attention, or what states to visit, facilitate adaptively expanding or reducing a state space? Under what circumstances (or perhaps unfortunate sequences of early observations; e.g., see (Smith et al., 2019a)) might an agent fail to sample locations in its environment that would have allowed for adaptive model expansions or reductions? How might the structural priors or hyperpriors inherited by an organism in a particular environmental niche guide adaptive action-guided structure learning? These questions touch on deep problems in learning while acting – here in the context of learning structure – that active inference models may be especially well equipped to address (i.e., due to the intrinsic information-seeking drive provided by expected free energy minimization; (Mirza et al., 2016; Parr and Friston, 2017; Tschantz et al., 2019)).

Here it is also again worth highlighting that, for agents acting within a particular environmental niche, a model’s accuracy need not correspond to a model’s adaptiveness (i.e., fitness or marginal likelihood). In some cases, it may be more adaptive for an agent to efficiently learn to model only the features in its environment relevant to its own survival, goal-achievement, etc. (Baltieri and Buckley, 2019; Tschantz et al., 2019), and organisms can benefit in some cases from acting based on false beliefs (McKay and Dennett, 2009); e.g., simplifications or heuristic priors (Al-Muhaideb and Menai, 2011; Gigerenzer and Gaissmaier, 2011). The distinction between accurate and adaptive models – where adaptive is operationally defined as free energy minimizing – is the complexity or KL-Divergence between posteriors and priors (i.e., not related to the notion of satisficing (Oaksford and Chater, 2003)). Interestingly, it is exactly this (complexity) term that structure learning by Bayesian model reduction optimizes – which, as discussed above, is also important to consider when interpreting the ‘accuracy’ of our model reduction results above across different simulated patterns of experience.

In conclusion, the Active Inference scheme we have described illustrates feature integration in the service of conceptual inference: it can successfully simulate simple forms of concept acquisition and concept differentiation (i.e. increasing granularity), and it spontaneously affords one-shot generalization.

Finally, it speaks to empirical work in which behavioral tasks could be designed to fit such models, which would allow investigation of individual differences in concept learning and its neural basis. For example, such a model can simulate (neuronal) belief updating to predict neuroimaging responses as we illustrated above; i.e., to identify the neural networks engaged in evidence accumulation and learning (Schwartenbeck et al., 2015). In principle, the model parameters (e.g., **A** matrix precision) can also be fit to behavioral choices and reaction times – and thereby phenotype subjects in terms of the priors under which they infer and learn (Schwartenbeck and Friston, 2016). This approach could therefore advance neurocomputational approaches to concept learning in several directions.

The issues raised here have a special importance in an active inference setting, in the sense that minimizing expected free energy means choosing policies that maximize information gain. In previous work, we have dealt with selection of policies that yield informative data to help infer states of the world, or to actively learn the parameters of the generative model. The results presented here propose an additional challenge for future work: how do we select policies such that we maximize information gain about the structure of the generative model itself?

Future research should investigate whether this basic approach can be successfully extended to meet this challenge and assess the additional benefits it might offer when applied to more complex real-world problems.

### Software note

Although the generative model – specified by the various matrices described in this paper – changes from application to application, the belief updates are generic and can be implemented using standard routines (here **spm_MDP_VB_X.m**). These routines are available as Matlab code in the DEM toolbox of the most recent version of SPM academic software: http://www.fil.ion.ucl.ac.uk/spm/. The simulations in this paper can be reproduced (and customized) via running the Matlab code included here is supplementary material (**Concepts_model.m**).

## Supporting information

Concepts_model.m

## Acknowledgment

This manuscript has been released as a Pre-Print at bioRxiv (Smith et al., 2019b).

## Footnotes

1 Generally motivated by starting with a finite parametric model and taking the limit as the number of latent states with more parameters tends to infinity.

2 However, “risky” reporting behavior could be elicited by manipulating the strengths of the agent’s preferences such that it placed a very high value on reporting specific categories correctly (i.e., relative to how much it disliked reporting incorrectly).

3 To break the symmetry of the uniform distribution, we added small amounts of Gaussian noise (with a variance of .001) to avoid getting stuck in local free energy minima – or any locations with zero free energy gradient – during learning (e.g., in this case, exactly equal concentration parameter updates could repeatedly occur across state-outcome mappings on each trial and prevent the agent from converging to a more accurate model).

